# Syntopic call diversity among morphologically similar calling morphs of the paleotropical katydid of genus *Mecopoda* (Tettigoniidae) from an Indo-Burma biodiversity hotspot

**DOI:** 10.1101/2024.11.20.624612

**Authors:** Aarini Ghosh, Vivek Dasoju, Jishnu Borgohain, Ashique Rahman, Bittu Kaveri Rajaraman

**Affiliations:** Department of Biology, Trivedi School of Biosciences, Ashoka University, Sonipat, Haryana, 131029, India; Department of Psychology, Ashoka University, Sonipat, Haryana, 131029, India; Department of Environmental Studies, Ashoka University, Sonipat, Haryana, 131029, India

**Keywords:** Calling song types, phylogenetic, acoustic diversity, bushcricket, North-East India, Garo hills

## Abstract

The acoustically diverse family of Tettigoniidae (Ensifera, Orthoptera) (commonly called bushcrickets or katydids), constitutes an excellent system to study the evolution of calling songs. *Mecopoda* is a genus that is particularly interesting because of the many morphologically cryptic calling morphs that have been described across South, East, and Southeast Asia, some of which are considered endemic to India. We describe five new syntopic and sympatric calling morphs of the genus *Mecopoda* that are morphologically similar but acoustically diverse from a sub-tropical forest in Meghalaya, in the Indo-Burma biodiversity hotspot. We report the morphological and acoustic characterization of these five call types, finding morphological overlap but well-separated acoustic clusters based on a PCA analysis of call temporal and spectral features. We also sequenced the COI gene mitochondrial gene of these five call types as well as other call types previously reported from India and performed a phylogenetic analysis of this gene relative to previously reported COI gene sequences from genus *Mecopoda.* Phylogenetic results suggest our five call types are not mutually monophyletic and relate from different subspecies of *Mecopoda*, of which three call types are genetically well separated from other calling morphs. This broadens our understanding of the evolution of acoustic diversity among paleotropical katydids.

## Introduction

Katydids or bushcrickets of the genus *Mecopoda* (superfamily Tettigoniidae, suborder Ensifera, order Orthoptera) are abundantly found in South Asia, East Asia, and Southeast Asia (Song *et al.,* 2015; Liu *et al*., 2020; Tiwari and Diwakar, 2020; Heller *et al.,* 2021). The first recorded description of *Mecopoda elongata* was made by Linnaeus (1758), who called it *Gryllus (Tettigonia) elongata.* This was later differently named as *M. maculata* by Serville, which constituted the type specimen for the genus *Mecopoda* (1831). A number of later studies characterized *Mecopoda* morphologically (Ingrisch & Shishodia., 2000; Ingrisch & Garai., 2001), with a focus on genital (Dutta *et al.,* 2018) and wing morphology (Heller *et al.,* 2021).

*Mecopoda,* like many other Ensiferans, produces mating advertisement calls (Strauß & Lakes-Harlan., 2013), with males typically rubbing together specialized forewings to stridulate (Greenfield., 2016). The left wing is equipped on the underside with a file, consisting of a series of cuticular protrusions known as pegs or teeth, which are rubbed against a hardened plectrum on the right wing during the opening and closing of the wings (Stumpner, 2013). Males of each species typically produce calls with specific temporal and spectral features. However various ecological pressures may select for mutations leading to changes in the spectral or temporal features of the call. Since females of most species have a preference for specific conspecific call characteristics, acoustic divergence can lead to assortative mating depending on the interaction between the range of male call types and female call preferences within a population. In the course of evolution, assortative mating can also lead to behavioral reproductive isolation, which in turn can enable morphological and genetic divergence between call types and even eventually speciation (Wilkins *et al.,* 2012; Gray & Cade., 2000).

The first bioacoustic characterization of *Mecopoda* songs was done by Sismondo (1990) who described various “song species”, many of which are morphologically cryptic (Liu *et al.,* 2020; Nityananda & Balakrishnan., 2006; Tiwari & Diwakar 2022). Morphological and geographic differentiation of these various call types has not been conclusive (Karny., 1924), with many superficial differences in coloration within each calling type (Heller *et al.,* 2021). This disjunction between morphology and bioacoustics has posed a challenge for species identification (Balakrishnan, 2005), with Otte (1992) positing that each call type is a separate species.

Whole genome data may help shed light on taxonomic resolution. So far, most work has combined single gene data with acoustic and morphological markers for a more robust categorization of the wide number of species and subgroups from regions of East and Southeast Asia within the genus *Mecopoda*. *Mecopoda elongata* is considered a superspecies, with a trilling supersubspecies with narrower wings called *Mecopoda elongata niponensis* and a chirping supersubspecies with broad tegmina and long files called *Mecopoda elongata confracta* (OSF., 2024). Some initial genetic data using the mitochondrial COI gene that codes for cytochrome c oxidase subunit I (Hebert *et al.,* 2009) and acoustic analysis has emerged for seemingly cryptic species in East Asia that produce more complicated songs with grouped chirps (Liu *et al.,* 2019), and some of these have been grouped as a subsuperspecies of *Mecopoda elongata* called *M. elongata minor*.

In the meantime, it is hard to conclusively define the number of species of this genus, but so far from the Indian subcontinent, only four have been identified at the species level in the literature: *Mecopoda elongata* consisting of various call types (Nityananda & Balakrishnan, 2006), *M. fallax* (Tiwari & Diwakar., 2022), *Mecopoda pallida* (Walker, 1869; Liu *et al.,* 2020), *Mecopoda sp* (Ashwathanaryana & Ashwath., 1994), along with various unplaced call types (Tiwari & Diwakar., 2022). A variety of call types of *M. elongata* have been described from India that have not been reported elsewhere and may be considered endemic to India (Heller *et al.,* 2021), with some of these calling types showing sympatry and even syntopy (Nityananda & Balakrishnan., 2006; Tiwari & Diwakar., 2022). In this paper, we have carried out morphological, acoustic, and genetic analysis of a set of five sympatric call types from the genus *Mecopoda* found in the tropical rainforests of Meghalaya, along with COI gene sequencing and phylogenetic analysis of these as well as the call types reported from South India (Nityananda & Balakrishnan., 2006).

## Material and Methods

### Study sites

Acoustic and specimen sampling was done inside and outside protected areas of Nokrek National Park located in Garo Hills of Meghalaya, India, with due permission. This protected area consists of sub-tropical, tropical evergreen, and tropical semi-evergreen forest types (Prabhu *et al.,* 2010). The peripheral Daribokgre village has tropical bushes, areca palm, and wild orange plantations (Fig 1B). Daribokgre village and Nokrek National Park are at an altitude of 500-1190m a.s.l., GPS location-25° 28’ 37.56’’ N; 90° 19’ 5.0628’’ E (Fig 1A). In the monsoon season (July-September) the precipitation rate was 500-2000 mm and the minimum and maximum temperatures throughout the year were 35°C and 10°C respectively (http://megagriculture.gov.in). Nokrek National Park falls within the Indo-Burma biodiversity hotspot (Venkataraman & Sivaperuman., 2018) and also shares a boundary with the Eastern Himalayan hotspot (Fig 1).

**Figure 1.**
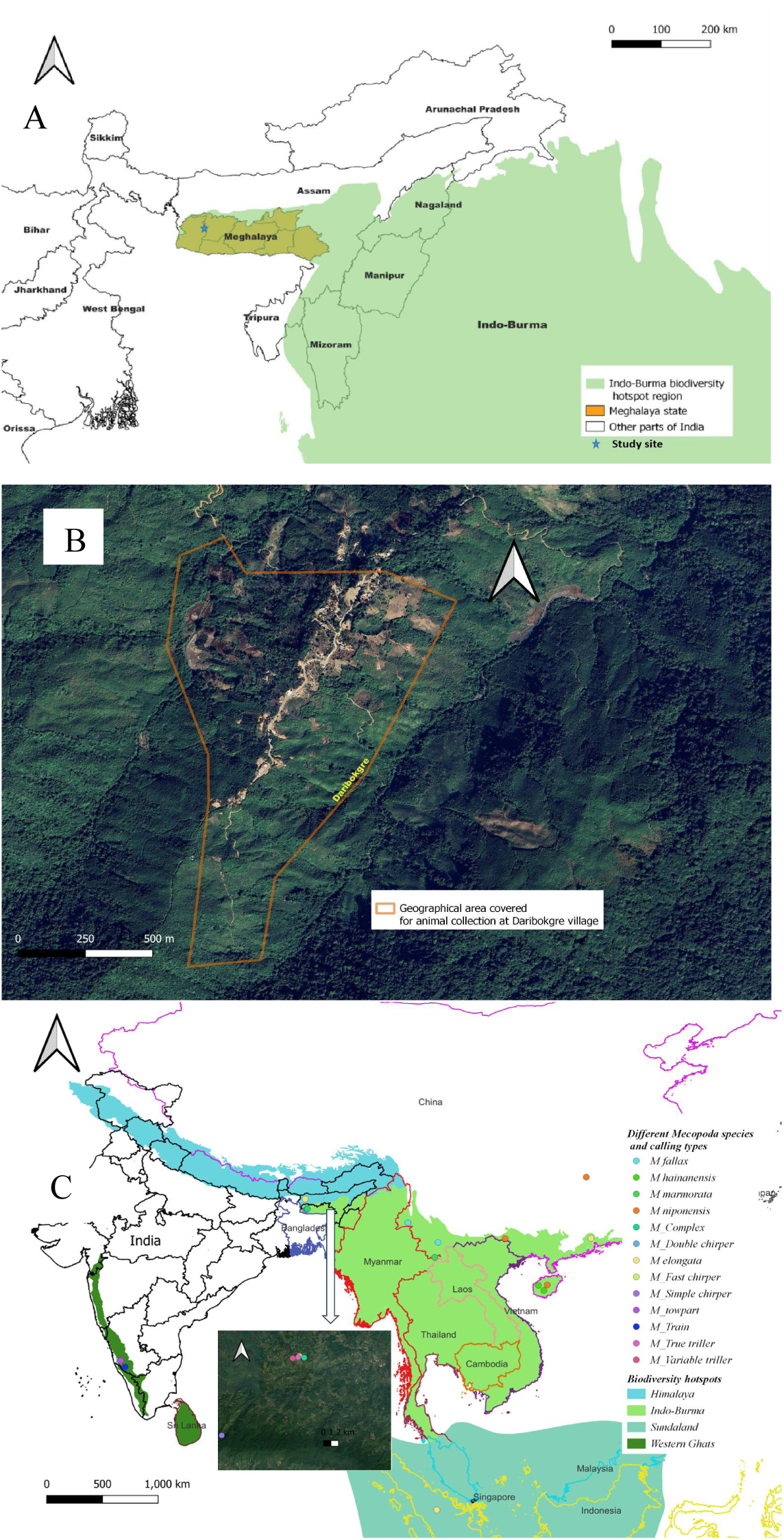
(A) Map showing the study site of specimen collection and boundary of Indo-Burma biodiversity hotspot with state boundary of Meghalaya districts in India. (B) Open street map showing the geographical area covered for specimen and sound collection at Daribokgere village, East Garo Hills in the periphery of Nokrek National Park. (C) Map of all *Mecopoda* COI gene sequence collection sites (data from Shen & He (2019) and our collection, throughout South and Southeast Asia). Different color shades represent different biodiversity hotspots.

### Specimen collection and preservation

*Mecopoda* are largely found on bushes, from lower to middle vertical strata. They inhabit a variety of habitats, from deciduous forests to thorny scrub areas, to dense sub-tropical forests. 20 male specimens were collected by localizing them from their calling positions in the bushes. Collections were done over the span of 4-5 days around the no-moon period of November of 2021 and 2022 between 7:00 pm - 10:00 midnight. All specimens for taxonomic identification were dry preserved. A few male specimens from each call type were preserved in 70% ethyl alcohol for morphometric measurements at the Neuroethology lab of Ashoka University.

### Acoustic recording and analyses

Calling songs of males of all five call types (n=6 each) were recorded in the wild using a Pettersson M500 USB ultrasound microphone mounted on a laptop (NOKIA) at a sampling rate of 500 kHz. The recording was done from a 5-foot distance from the caller to avoid clipping. A digital hygrometer (STORE99 WHDZ HTC-1) was used to measure the ambient temperature and humidity each day and found to vary very little over the recording span of 4-5 days when there was low or no moon in November of 2021 and 2022 (average temperature was 21°C +/− 0.5 degree and average humidity 84% +/−3, median +/− interquartile range). Temporal and spectral parameters of acoustic recordings were analyzed using Bat Sound 3.31 and Audacity software Version 3.0.5, for 10-20 syllables per call. Power spectrum analysis and spectrograms were made in Bat Sound standard sound analysis software 3.31.

We describe the temporal structure of the call (Fig. 2) with the following definitions similar to those used in the literature (Tiwari & Diwakar 2022; Heller et al., 2021; Baker & Chesmore, 2020). One **syllable** corresponds to the sound pulse in an oscillogram caused by a single strike of the forewings against each other, and **syllable duration** is calculated as the time period spanning that pulse. **Inter-syllable duration** is the time interval between two syllables, and a **syllable period** is the period from the beginning of one pulse to the next, the sum of syllable duration and inter-syllable duration. The **syllable repetition rate** is calculated as a reciprocal of the syllable period. An unbroken long series of syllables is called a trill; but if syllables show further grouping and pauses, the smallest first-order grouping of syllables that repeats itself is called an **echeme or chirp**, characterized by an **echeme duration and inter-echeme duration. The echeme period** is the sum of the echeme duration and time between two echemes (Fig 2). Syllable variations across echemes, when present, were also noted down.

**Figure 2.**
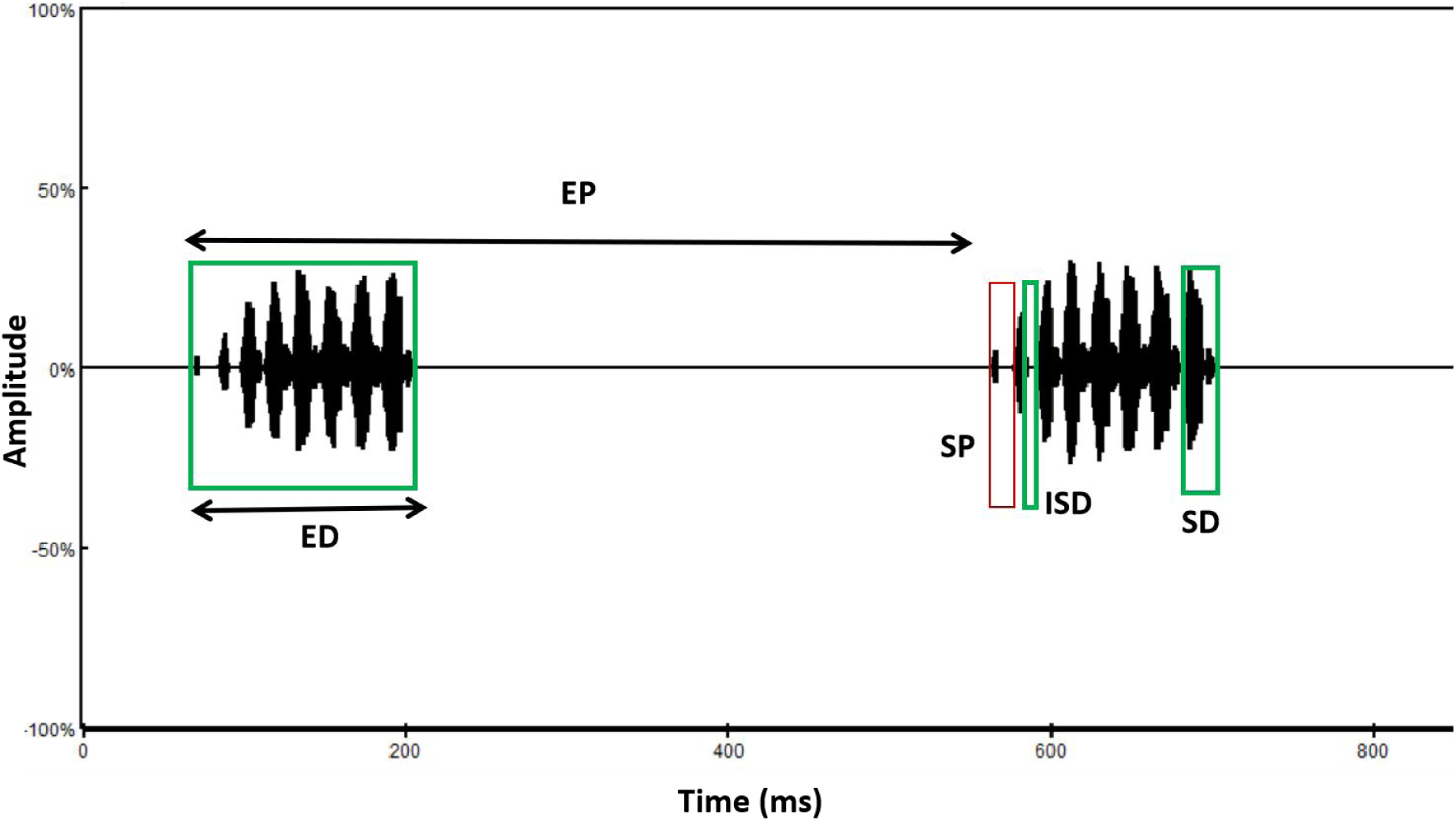
Schematic representation of an oscillogram of a katydid song denoting all temporal parameters using the following definition by Heller *et al*., 2021.

**Abbreviations**

**SD-** Syllable Duration

**ISD-** Inter-Syllable Duration

**SP-** Syllable Period

**EP-** Echeme Period

**SyRR-** Syllable Repetition Rate

**S/E-** Syllables in each Echeme

**PF-** Peak Frequency

**FF-** Fundamental Frequency

**BW-** Bandwidth

To characterize spectral features, we noted down the peak frequency, fundamental frequency, and bandwidth by analyzing the power spectrum of each call. **Fundamental frequency** was defined as the first peak and **peak frequency** as the highest peak in the power spectrum (Baker & Chesmore 2020). **Bandwidth** was calculated as the difference between minimum and maximum frequency −20dB below the peak frequency (Heller *et al*., 2021; Tiwari & Diwakar., 2022).

### Morphological structures and measurements

Morphological structures were studied under a stereo zoom microscope, Leica S9i (Leica Microsystems, Germany). Measurements of 21 quantitative morphological characters and a full qualitative study of 20 males (4 individuals from each of 5 call types) were made using the analysis tool of Leica Application Suite version LAS V4.13.0 installed on a computer running a Windows 10 Operating system in the Department of Biology, Ashoka University, Haryana. To measure full body length for calibration, we used Vernier Calipers.

### Imaging

Scanning Electron microscope imaging of the stridulatory teeth on the wings was done at 1–2 kV resolution using JSM-6100 (JEOL, Japan) in the Department of Sophisticated Analytical Instrumentation Facility, CIL and UCIM, Panjab University, Chandigarh, India.

### Statistical analysis

We performed a Shapiro-Wilk test on the acoustic data to check the normality of the data set. We used 10 of each type of acoustic unit (syllables, echemes, etc.) for analysis from each individual’s call, and these data were normality distributed; however, the data across individuals do not show a normal distribution. We therefore used non-parametric tests and represented the data using medians and inter-quartile range (IQR). To check for differences across all 5 call types together we ran a Kruskal-Walli’s test, and Dunn’s test was used for pairwise comparisons between call types. All statistical tests were performed in RStudio software. For multidimensional analysis of acoustic and morphological data, PCA (principal component analysis) was performed for dimensionality reduction to visualize data clusters in RStudio. The hierarchical clustering of traits was done with acoustic data and morphological data using the hierarchical clustering method in RStudio.

### Phylogenetic analysis

Single samples of calling males were collected from Meghalaya and Western Ghats and preserved in absolute alcohol. Most were sent for COI gene (a 250 bp fragment) Sanger sequencing at a facility run by Medauxin (http://www.medauxin.com/). For one call type, “Simple Chirper”, the COI gene sequence was retrieved from a whole mitochondrial genome analysis by shotgun sequencing with an overlap of 40% conducted by MolSys Pvt. Ltd.We used the software MITOS to do the mitochondrial genome assemblage.

We cut all our COI sequences to select high-quality portions, and then performed multiple alignments of our sequences along with published COI data for *Mecopoda* from NCBI (see the full list in Table 9), using *Ducetia japonica* as the outgroup. We then constructed a Maximum likelihood phylogenetic tree using MEGA11. We also mapped the geographic locations from which all the published and newly sequenced COI gene data of different species and acoustic morphs of *Mecopoda* were collected from around the world (Fig. 1C).

## Results

### Distribution of Animals

The five different call types of *Mecopoda elongata* we found were collected from one location at Nokrek National Park in West Garo Hills of Meghalaya. Of these call types, we also found the one we call the “Complex” call type in many other locations of Meghalaya, including the Khasi hills. The vegetation of Nokrek National Park is mostly evergreen and mixed forest with patches of cultivation area (Fig 3). Calling males were mostly found in bushes and midstory vegetation. Around Nokrek, all five call types are syntopic and can be found co-occurring and calling at the same time of the day in the same microhabitat.

**Figure 3.**
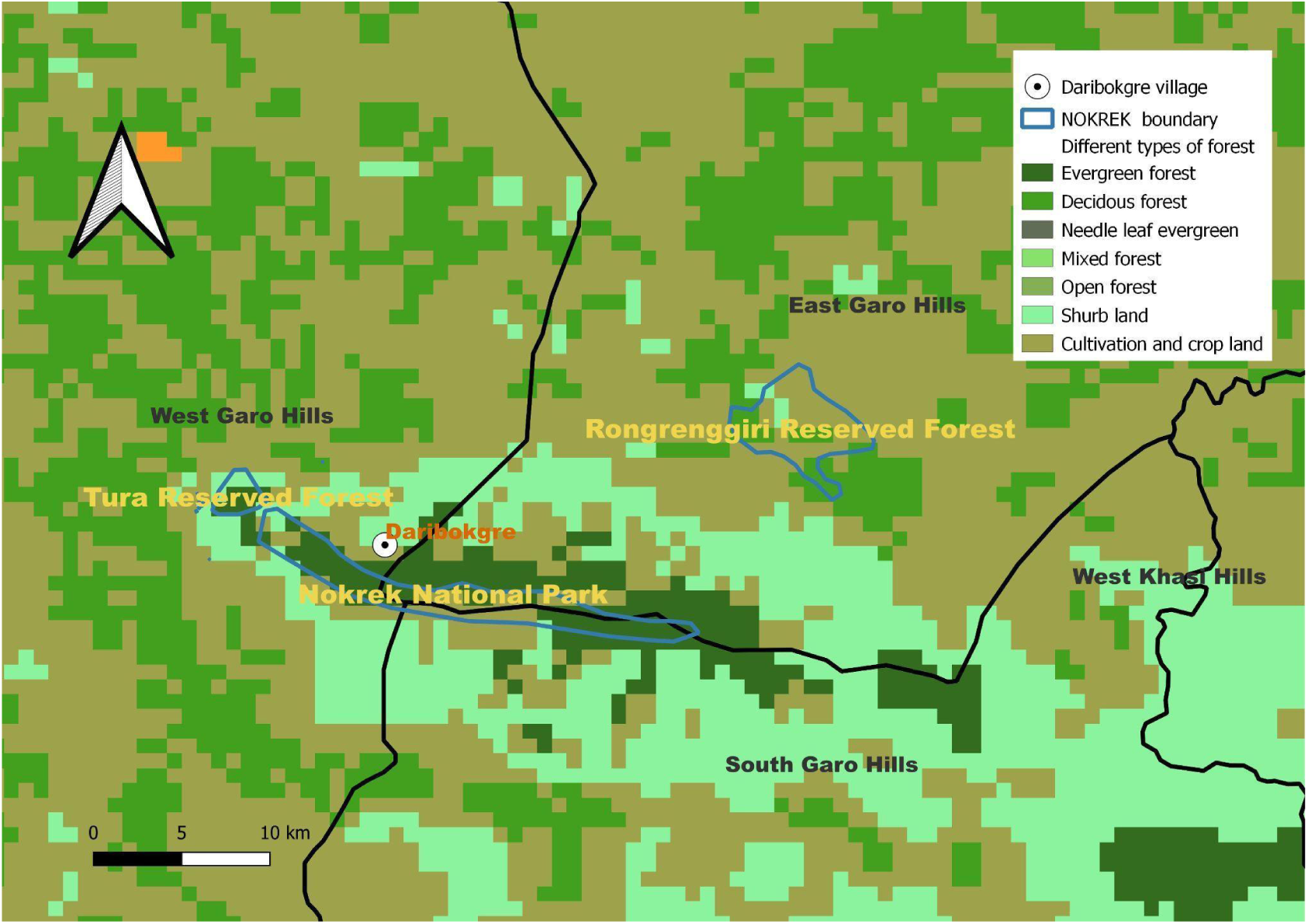
LULC map showing different vegetation types at the core area of Nokrek National Park and its periphery.

**Abbreviations and call type summary:**

Mecopoda “Simple Chirper”: CH

Mecopoda “Fast Chirper”: FC

Mecopoda “True Triller”: TT

Mecopoda “Variable Triller”: VT

Mecopoda “Complex”: CO

### Acoustic analysis

The five call types of *Mecopoda* from the Garo Hills produce distinct calling songs. Detailed spectral and temporal analyses of each call type are described below.

#### 1. Simple Chirper call type

This is the simplest call type named “Simple Chirper”, with repeated distinct chirps with regular pauses (Fig 4). This call is very similar to that described for chirpers in South India (Nityananda & Balakrishnan, 2006). The echeme period is 0.65 +/− 0.15 s (median +/− IQR, n = 60 chirps). Each echeme consists of 8-10 syllables and each syllable has a syllable period of 0.043 +/− 0.01 s (median +/− IQR, n = 60 syllables) with a duration of 0.017 +/− 0.011 s (median +/− IQR, n = 60) and a median inter-syllable duration of 0.028 +/− 0.007 s (median +/− IQR, n = 60). There is some amplitude modulation among syllables in each chirp. All syllables have broadband spectral structure with a bandwidth of median 55.1 +/− 4.7 kHz, fundamental frequency of 8.9 +/− 1.6 kHz and peak frequency of 23 +/− 0.9 kHz (all median +/− IQR, n = 6 individuals).

**Figure 4.**
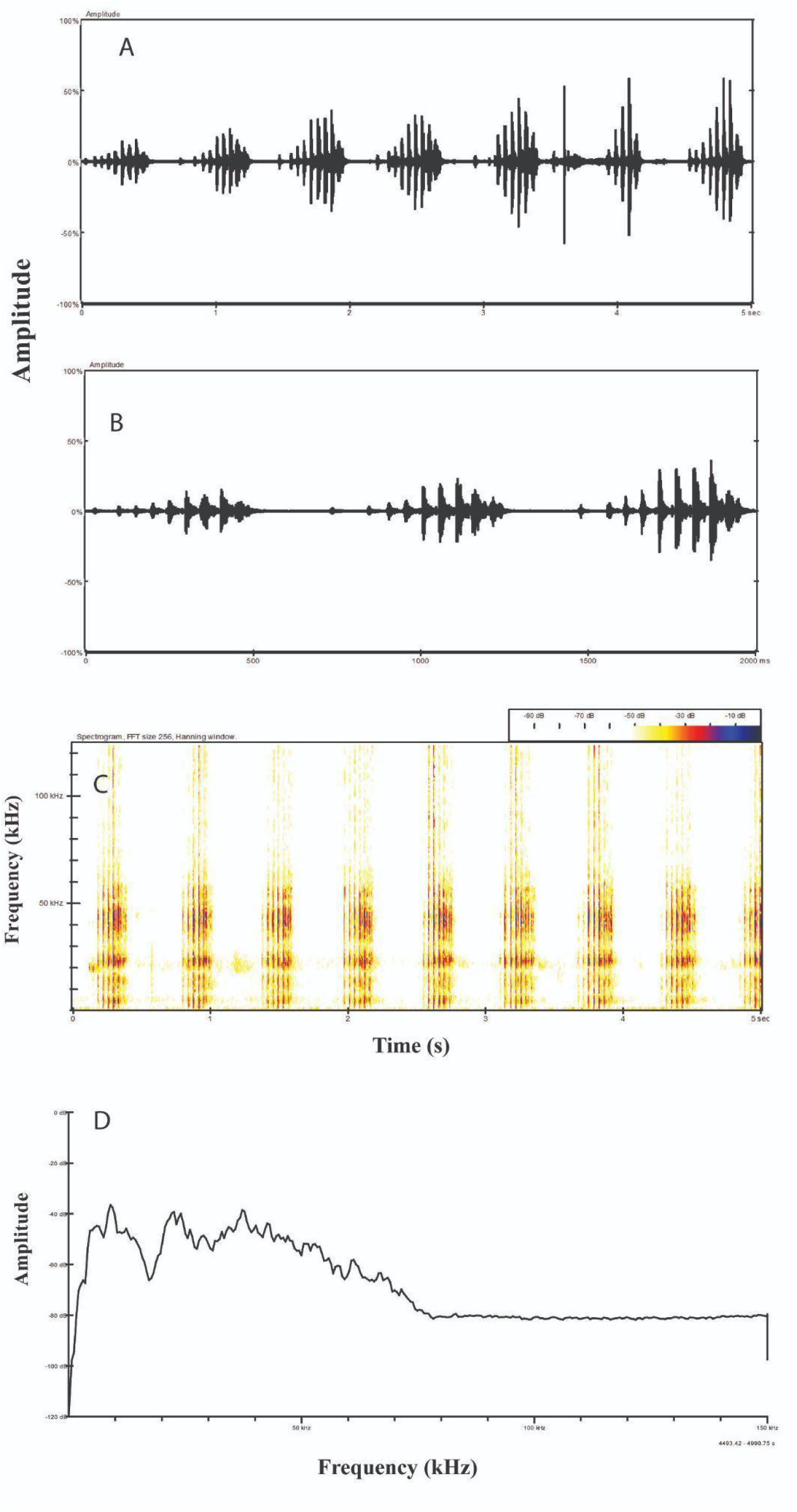
Temporal and spectral pattern with the power spectrum of a representative *Mecopoda* “Simple Chirper” call. (A-B) Oscillograms of calling songs at different time scales. (C) Spectrogram of the call. (D) Power spectrum of the call.

#### 2. Fast Chirper call type

This call type sounds like fast chirps interspersed with distinct pause, and hence we named it “Fast Chirper” (Fig 5). However, within each unit of sound corresponding to a wing stroke (Video link: https://doi.org/10.5281/zenodo.14053596) there is only one distinct syllable repeating with a unit period of 0.14 +/− 0.02 s (median +/− IQR, n = 80 syllables) and such a call would technically constitute a trill (Baker & Chesmore., 2020). The syllable duration is 0.097 +/− 0.029 s (median +/− IQR, n = 80 syllables) with an inter-syllable duration of 0.43 +/− 0.01 s (median +/− IQR, n = 80 syllables). This is a broadband call with a fundamental frequency of 7.7 +/− 1.8 kHz (median +/− IQR, n = 8 individuals) and peak frequency of 22.35 +/− 1.58 kHz (median +/− IQR, n = 8) with a bandwidth of 31.05 +/− 9.15 kHz (median +/− IQR, n = 8).

**Figure 5.**
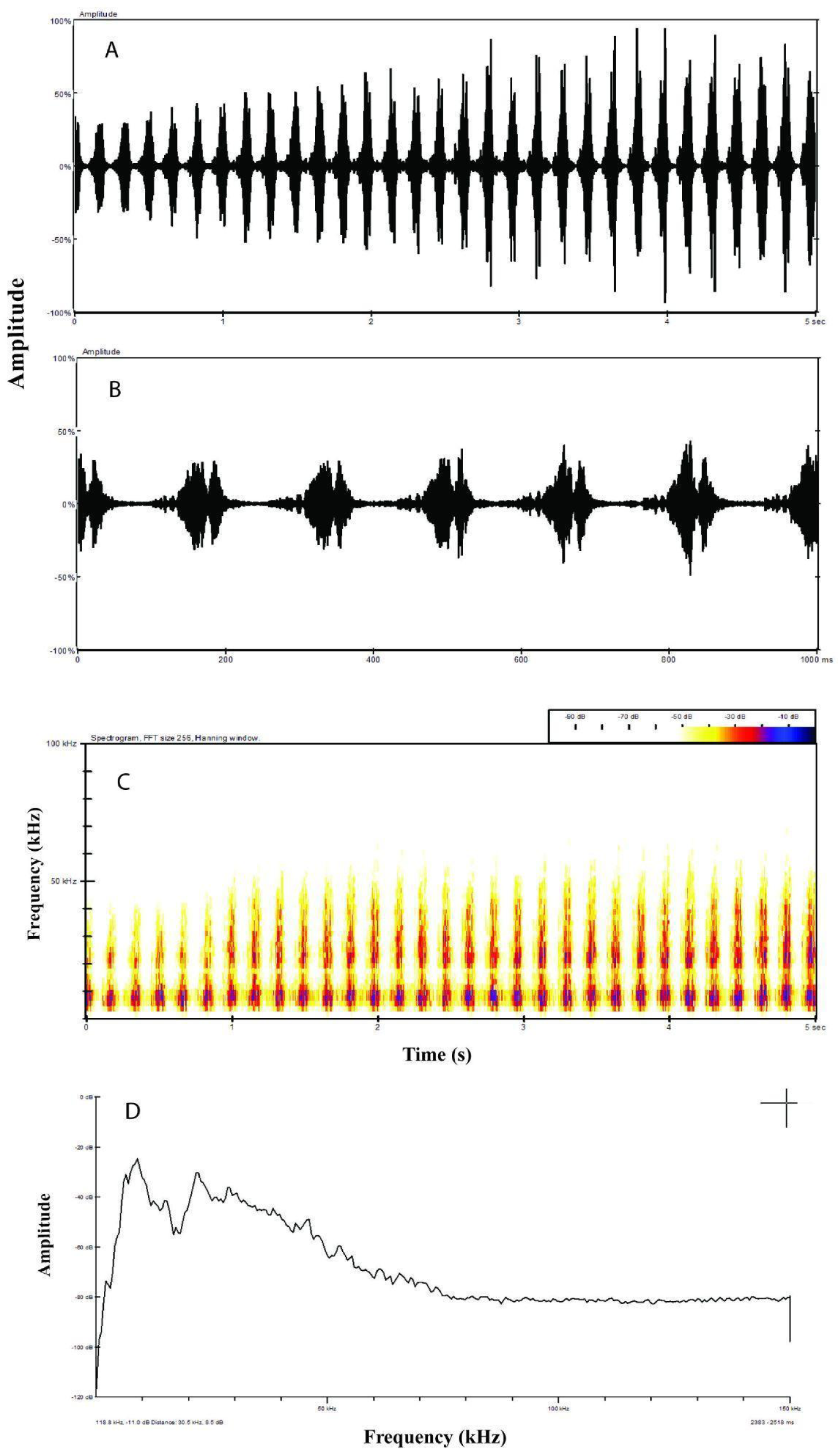
Temporal and spectral pattern with the power spectrum of a representative *Mecopoda* “Fast Chirper” call. (A-B) Oscillograms of calling songs in different time scales. (C) Spectrogram of the call. (D) Power spectrum of the call.

#### 3. True Triller call type

This call type is a trill with slight amplitude modulation (1.76, +/− 0.4 s, median +/− IQR, n = 33) which contains a continuous series of syllables without any distinct inter-syllable interval, where the repeated syllable segment (with two amplitude peaks) lasts for a duration of 0.045 +/− 0.006 s (median +/− IQR, n = 60 syllables) (Fig 6). We named this *Mecopoda* call type “True Triller”. This broadband call type has a fundamental frequency of 8.7 +/− 0.4 kHz (median +/− IQR, n = 6 individuals) and a peak frequency of 22.6 +/− 0.25 kHz (median +/− IQR, n=6), with a bandwidth of 42 +/− 10.2 kHz (median +/− IQR, n=6).

**Figure 6.**
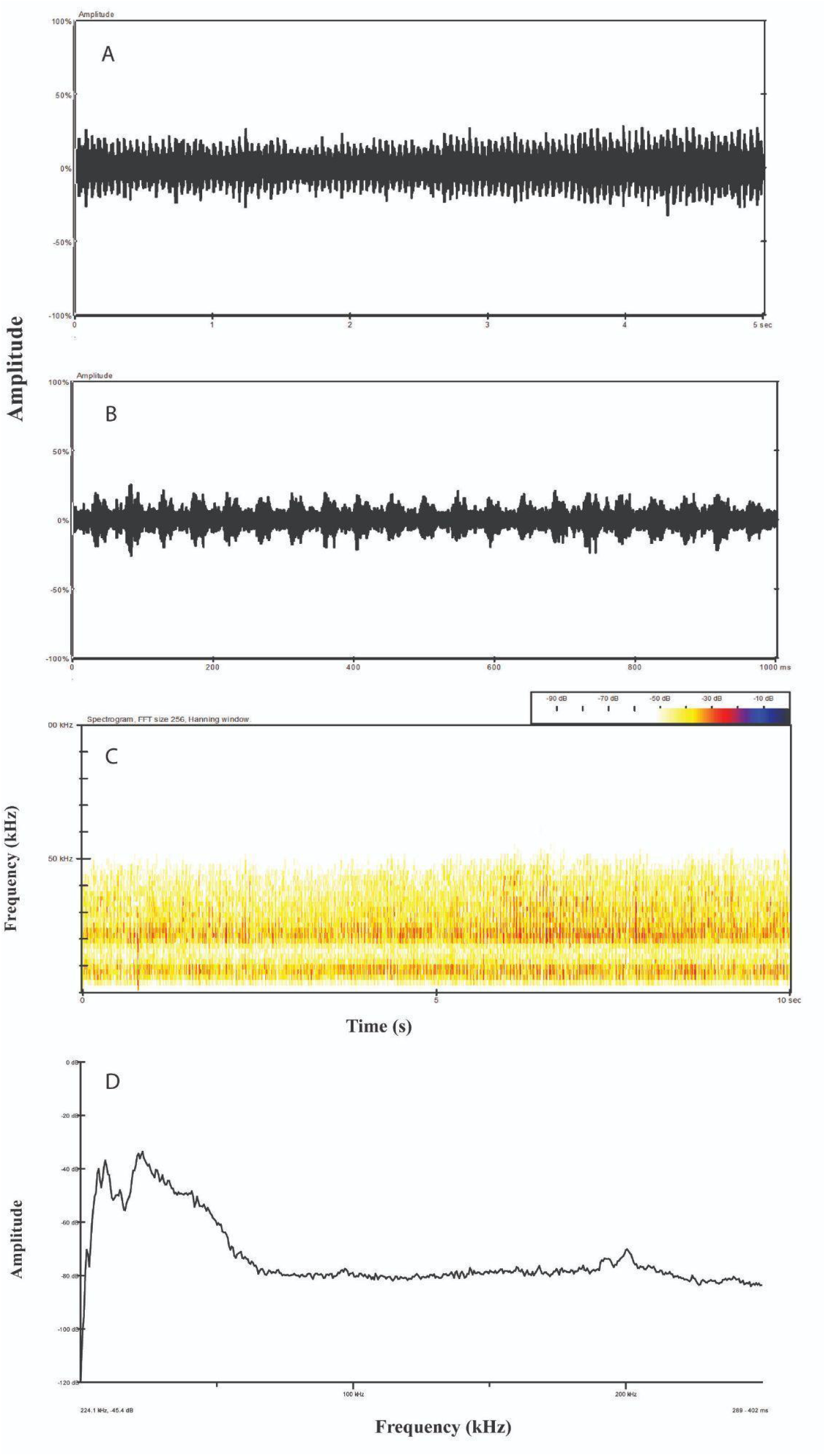
Temporal and spectral pattern of a representative *Mecopoda* “True Triller” call. (A-B) Oscillograms of its calling song at different time scales. (C) Spectrogram and (D) Power spectrum of the call.

#### 4. Variable Triller call type

The “Variable Triller” call type also has a continuous trill, but with strong amplitude variation, alternating between two different groups of syllables (Fig. 7). One group of syllables is low in amplitude and longer in duration (VT_II) while the other group is of shorter duration and ramps up to and from a higher amplitude (VT_I). There is no distinct inter-echeme interval between consecutive cycles of VT_I and VT_II. The syllable period is 0.053 +/− 0.008 s (median +/− IQR, n = 60 syllables) for the high-amplitude syllables and 0.083 +/− 0.017s (median +/− IQR, n = 60) for the low-amplitude syllables. The transition between the two-syllable sets is gradual: the short amplitude syllable segment consists of syllables of relatively mutually similar amplitude and syllable structure, followed by a ramping up to higher amplitude faster syllables followed by a decline back to the short amplitude syllables. While there is amplitude modification, there is no modification in frequency across the call, all syllables are high-frequency and broadband. The fundamental and peak frequencies are with 6.8 +/− 0.75 kHz and 23.4 +/− 1.5 kHz (median +/− IQR, n=11 animals) respectively, with a bandwidth of 45 +/− 8.85 (median +/− IQR, n=11).

**Figure 7.**
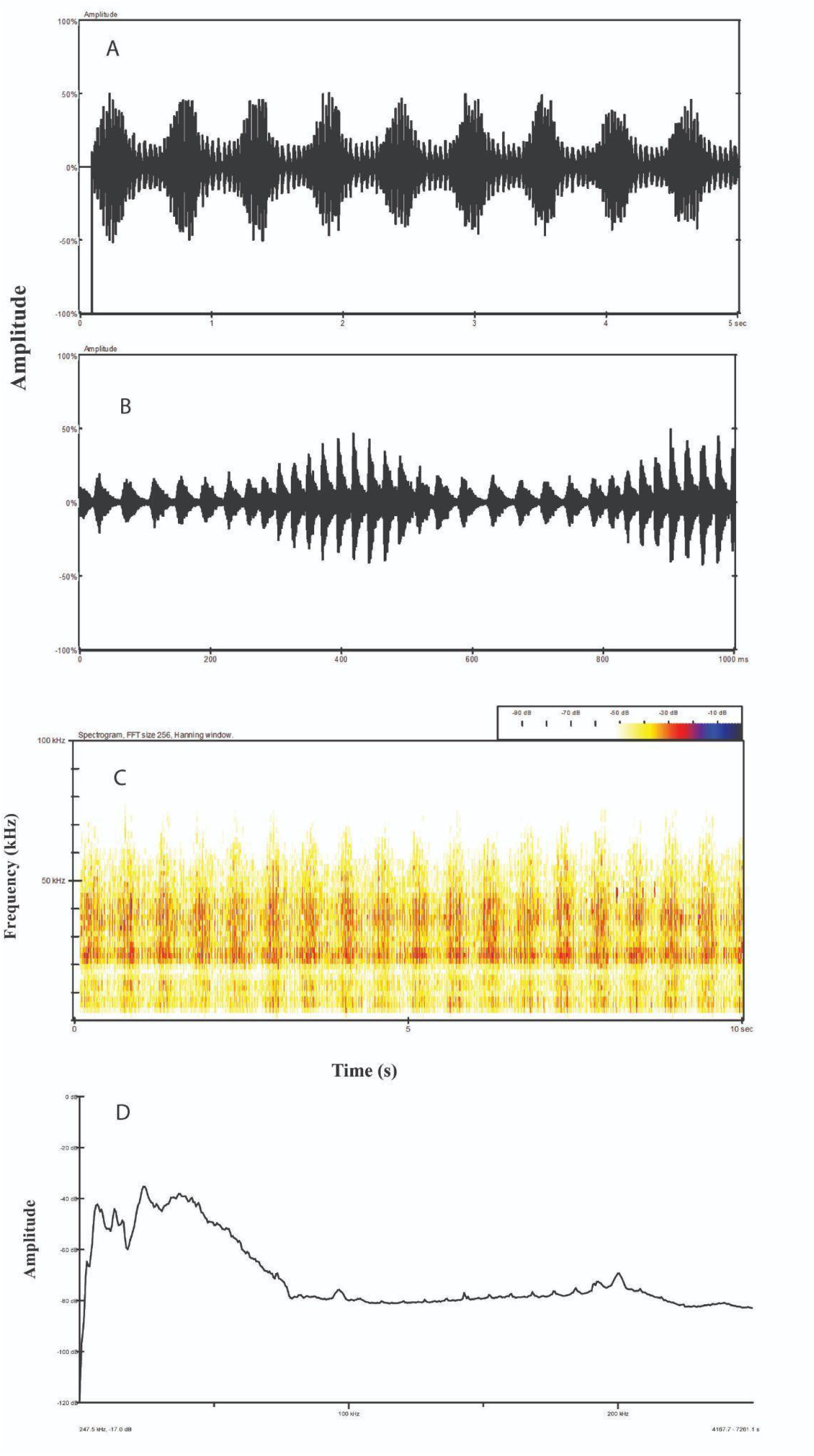
Temporal and spectral pattern of a representative *Mecopoda* “Variable Triller” call. Oscillogram of the calling song over (A) a 5 and (B) a 1 second timescale enabling a view of the syllable types present in the call (C) Spectrogram and (D) Power spectrum of the call.

**Figure 8.**
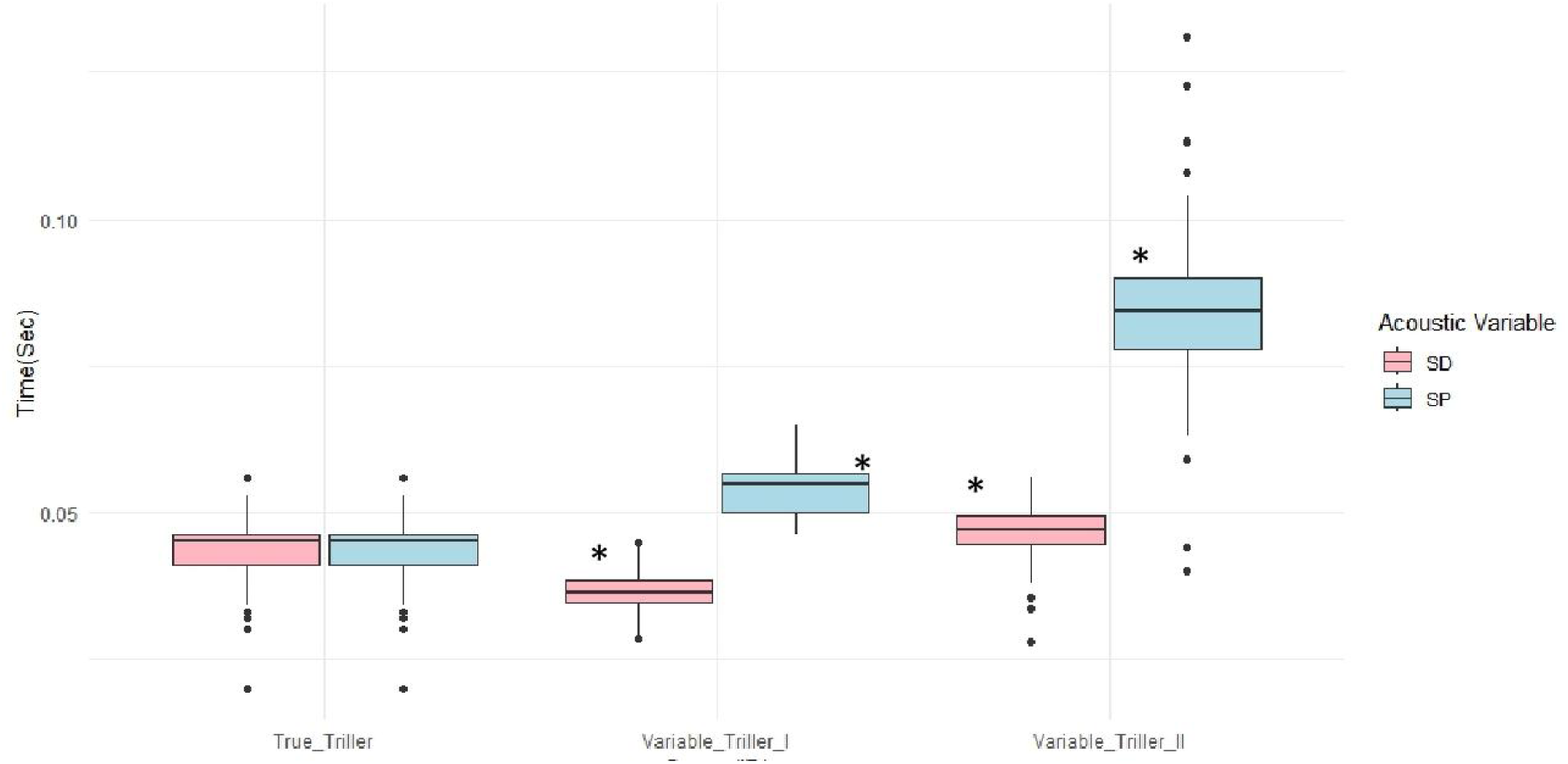
Temporal feature comparison between the “True Triller” call and the two parts of the “Variable Triller” call - the low amplitude longer duration portion (VT_II) versus the higher amplitude and shorter duration syllabi (VT_I).

### Comparison between the two triller call types

The main psychoacoustic difference between the two triller types is that “True Triller” sounds like a continuous trill without any variation in amplitude or syllable structure over time, but “Variable Triller” sounds like a trill with cyclical amplitude and syllable structure variation. There is significant difference in the syllable duration between the “True Triller” and both of the two-syllable types of “Variable Triller” (P = 0.0001 and 0.001 respectively for VT_I and VT_II). Also, the syllable period of VT_II, the low amplitude trill of “Variable Triller” is significantly longer than the syllable period of “True Triller” (P = 0.008), because of a longer inter-syllable duration in VT_II; “True Triller” has an indistinguishable inter-syllable duration from VT_I. The high amplitude portion of “Variable Triller”, VT_I also have a smaller inter-syllable duration and a statistically significantly smaller syllable duration than VT_II (P = 0.0003).

#### 5. Complex call type

This call type is called “Complex” because it has a complicated call structure, consisting of repeated, distinct, verses. (Fig 9). The average duration for the verse is 67.3 +/− 6.3s (median +/−IQR, n = 4 individuals). The first part of the verse is unique to the beginning of the call - it consists of stereotypical chirps, or echemes with regular pauses (CO_I). The echeme duration is 0.148 +/− 0.13 s (median +/− IQR, n = 60 echemes). Each echeme consists of 4-7 distinct syllables, with an echeme period of 0.3 +/− 0.47 s (median +/− IQR, n = 60 echemes). The syllable duration and syllable period are 0.019 +/− 0.012 s and 0.027 +/− 0.014 s (median +/− IQR, n=60 syllables) respectively. In the beginning, echemes are of low amplitude, and the amplitude shows a gradual increase followed by a gradual decrease before the beginning of the second part of the call. The second part of the verse (CO_II) consists of continuous indistinct fast chirps, where all syllables are higher in amplitude than in the first part of the call. The echeme structure of the second part of the verse varies from consisting of 4-9 syllables/echeme (median 5 +/− 1, n = 60 echemes), and therefore the echeme period is 0.276 +/− 0.128 s (median +/− IQR, n = 60). The syllable duration and syllable period are 0.023 +/− 0.017 s and 0.04 +/− 0.023 s (median +/− IQR, n=6) respectively. The third and last part of the verse is a trill consisting of indistinct syllables - without either any distinct pause between syllables or any grouping of syllables (CO_III). The syllable period is 0.035 +/− 0.018 s (median +/− IQR, n = 60 syllables). The average duration of the end part trill is 9 +/− 10 s (median +/− IQR, n = 6). After the third part of the verse ends, the animal repeats the verse cyclically. The fundamental and peak frequencies are 6.2 +/− 0.45 kHz (median +/− IQR, n=6) and 34.6 +/− 4.3 kHz (median +/− IQR, n=6) respectively, with a bandwidth of 56 +/− 6.95 (median +/− IQR, n=6).

**Figure 9.**
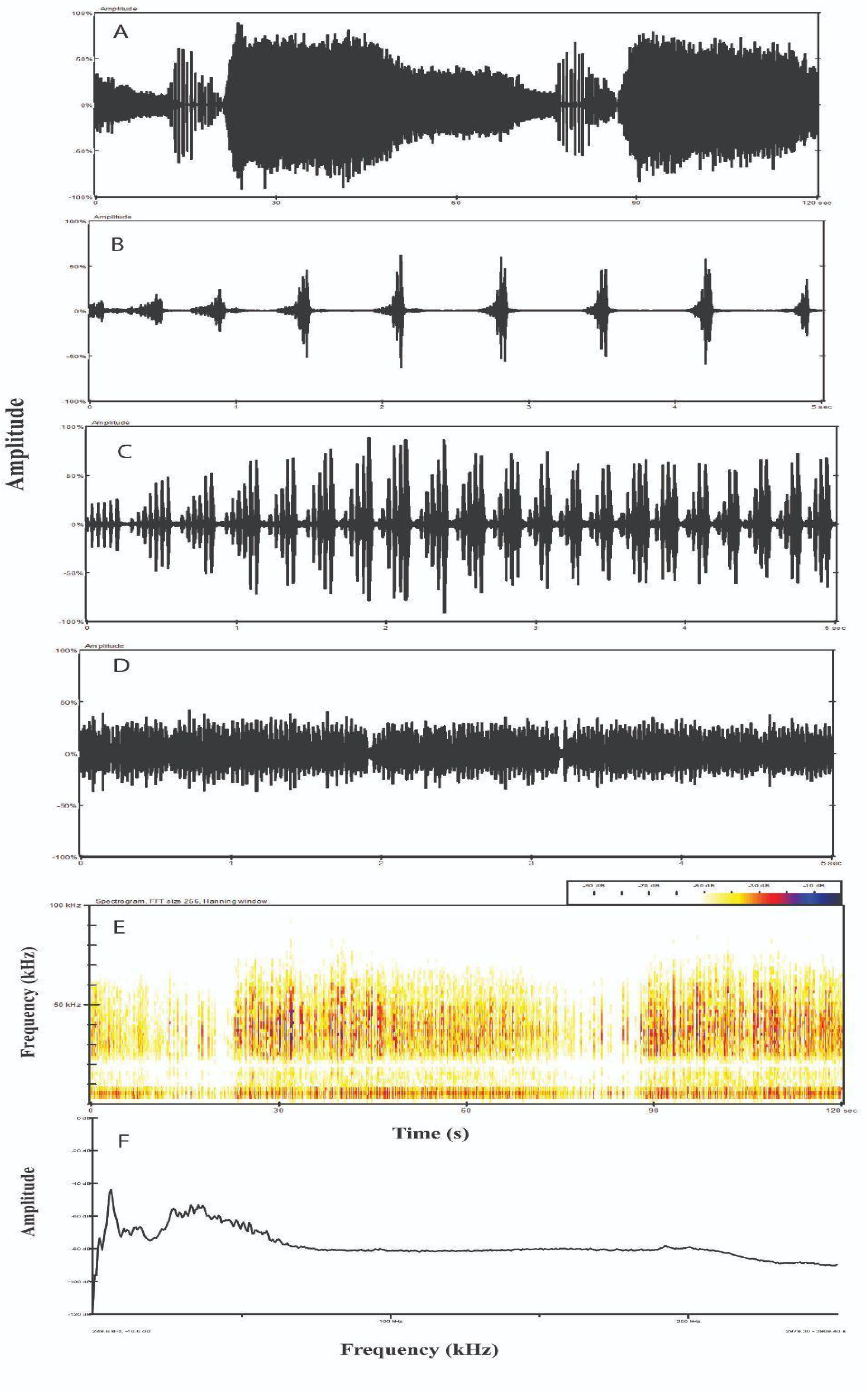
Temporal and spectral pattern of the power spectrum of a representative *Mecopoda* “Complex” call. Oscillogram of the calling song showing (A) the whole verse and its repeating structure, (B) the initial part of the call consisting of chirps, (C) the second part with indistinct chirps and (D) the ending trill consisting of indistinct syllables. (E) The spectrogram and (F) Power spectrum of the call.

### Comparison between different parts of Complex call

We found a statistically significant difference between the echeme structure of the different parts of the verse in the “Complex” call type, as well as in the syllable structure. Both the syllable duration and syllable period vary significantly between the three sections of the verse. CO_I and CO_III have significantly different syllable periods from CO_II (P = 0.0001 and 0.07), and there is substantial variation in the syllable duration in all sections of the verse. CO_III has significantly different syllable duration from CO_I and CO_II (P= 0.0001 and 0.003). The ending trill (CO_III) does not have a distinct inter-syllable period which means the syllable period is the same duration as the syllable duration (Fig 10A). There is a significant difference between the first two parts of the call with respect to the echeme period (P= 0.0037) (Fig 10B).

**Figure 10.**
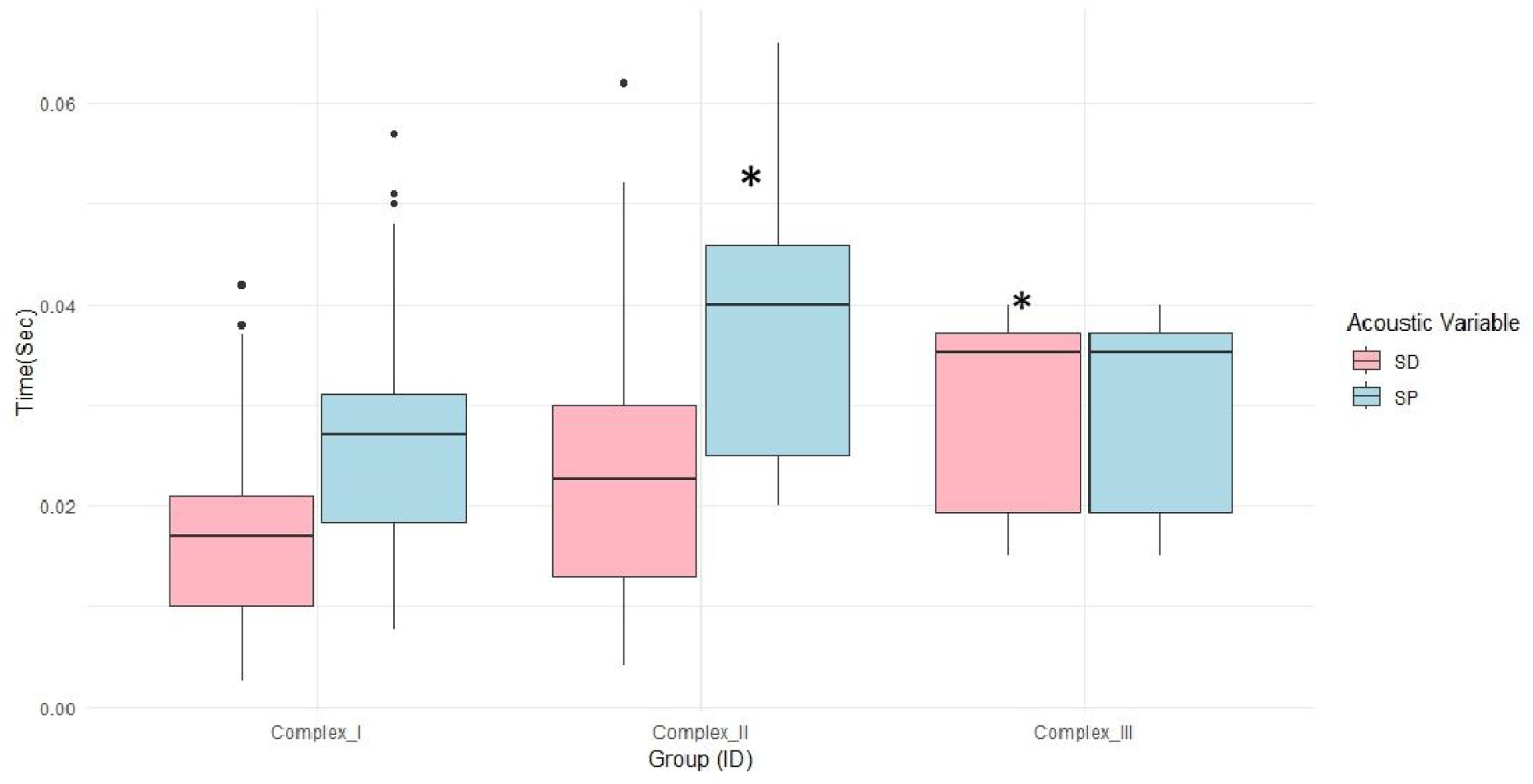
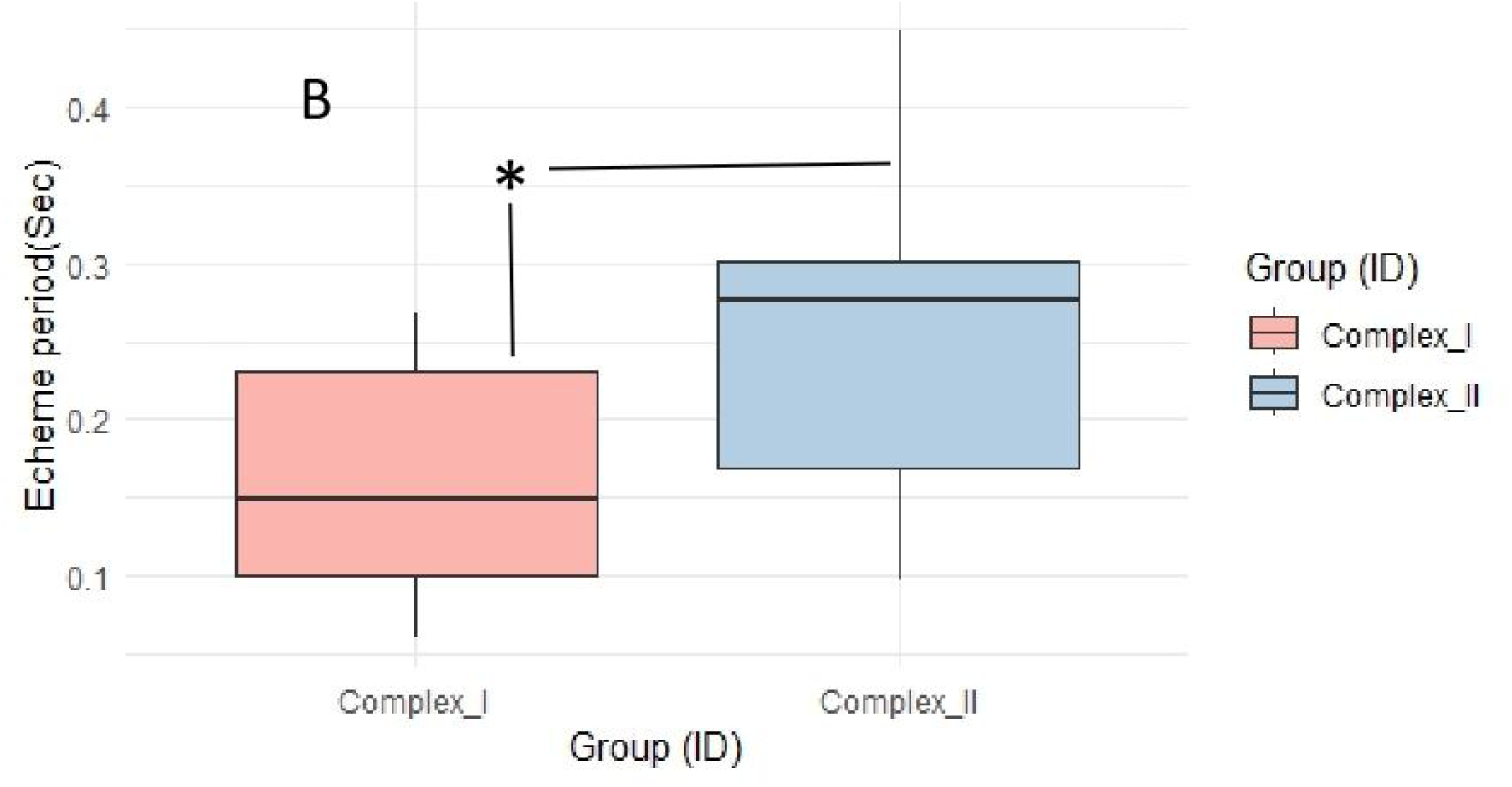
The difference in temporal features between different parts of the verse in the “Complex” call of *Mecopoda* call type. (A) Syllable features, and (B) Echeme features.

### Comparison of Different Parts of Complex Calls with Other Callers

Different parts of the “Complex” call have temporal characteristics which resemble parts of the call from other call types of *Mecopoda* such as “Simple Chirper”, “True Triller” and “Variable Triller”. The following bar plots were made to compare each part of the “Complex” call with other syntopic *Mecopoda* call types.

#### a. The difference between the starting chirping part of the “Complex” call and the “Simple Chirper” call

The temporal structure of “Simple Chirper” is somewhat different from the temporal structure of initial chirps of “Complex” call types (Fig 11B). The echeme period in the initial chirping part of the verse of “Complex” (CO_I) varies widely across individuals spanning the more restricted range of echeme periods for the chirps of “Simple Chirper” (P = 0.001; highly significant difference). The syllable period is significantly longer in the case of “Simple Chirper” than for CO_I, the initial chirps of “Complex” (P = 0.003) with no significant difference in syllable duration (P = 0.07) (Fig 11A).

**Figure 11.**
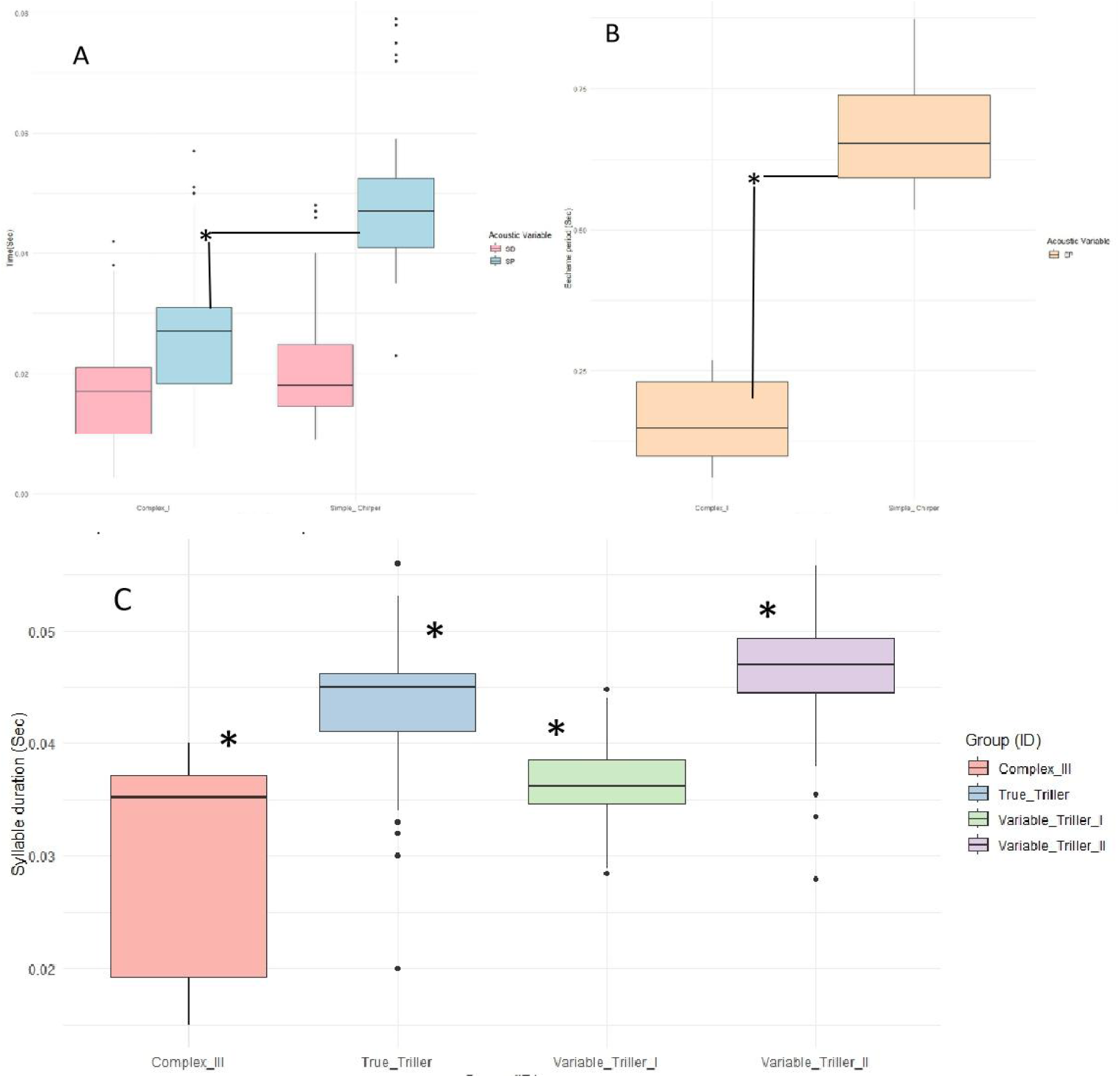
Box plot showing the temporal features difference between the initial chirping part of the verse of the “Complex” call and the call of “Simple Chirper” *Mecopoda* call types. (A) Echeme period (EP). (B) Syllable features: syllable duration (SD) and syllable period (SP). (C) Box plot showing the temporal feature differences between the third trilling section of the “Complex” call (CO_III) and the trilling calls of the “Variable Triller” and “True Triller” call types of *Mecopoda.* Note that there is no inter-syllable duration shown for trills such as that of TT and CO_III which have indistinct syllables.

#### b. End triller part of “Complex” call with “True Triller” and “Variable Triller”

The third part of the verse of “Complex” is shorter than “True Triller” (P=0.002) in that it consists of indistinct syllables, but the syllable duration of the trills of this third part of the verse is significantly shorter than the syllables of the “True Triller” call type. It is also significantly shorter than the syllables in the low amplitude syllables of “Variable Triller” and the high amplitude syllables of “Variable Triller” both (P= 0.009 and 0.0001) (Fig 11C).

In other words, the call portions of the complex call may look like a mixture of chirping and trilling call types, but the syllable structure of each of the chirping and trilling portions diverges from that of the chirper and triller call types.

### Acoustic clustering of call types

We conducted a principal component analysis (PCA) to visualize the separation of call types based on nine parameters calculable from the call data, both temporal and spectral (Table 1). The dimensional plot against both PC1 and PC2 shows the acoustic separation between the individuals of all five sympatric call types of *Mecopoda* at Nokrek National Park (Fig 12). The spread of the “Complex” call type is the widest and has three distinct clusters corresponding to the 3 different verse parts in its call, all of which are relatively well separated from other call types, especially the ending trill portion. “Simple chirper” and “Fast chirper” form tight clusters that are well separated from all the other callers in acoustic space. As seen in the direct comparison between triller calls, the higher amplitude syllables of “Variable Triller” VT_I overlap with the “True Triller” cluster.

**Figure 12.**
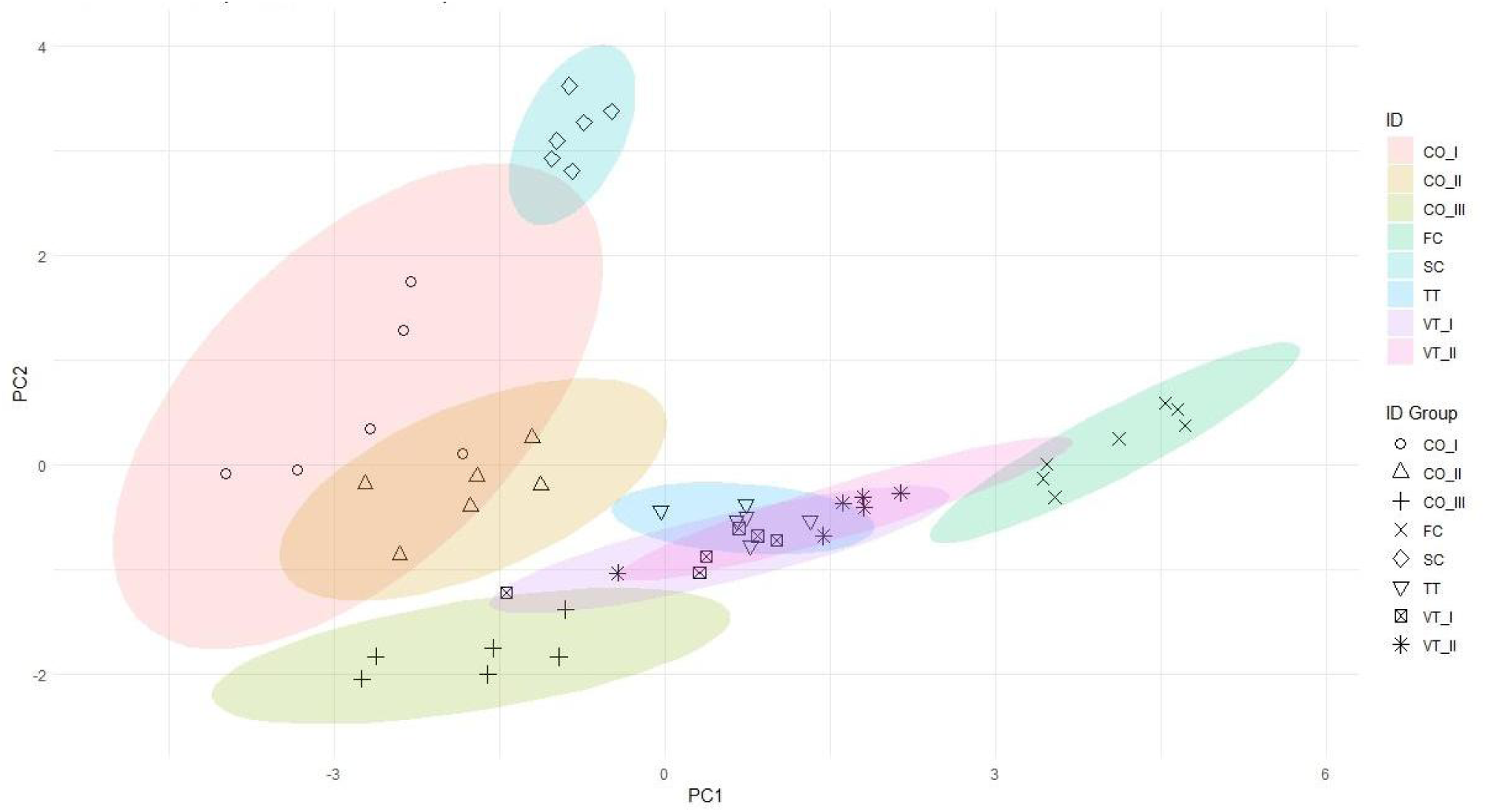
PCA visualizing the separation of call types based on temporal and spectral acoustic features. Each marker represents one individual, while colors denote the spread of call type data.

**Table 1.**
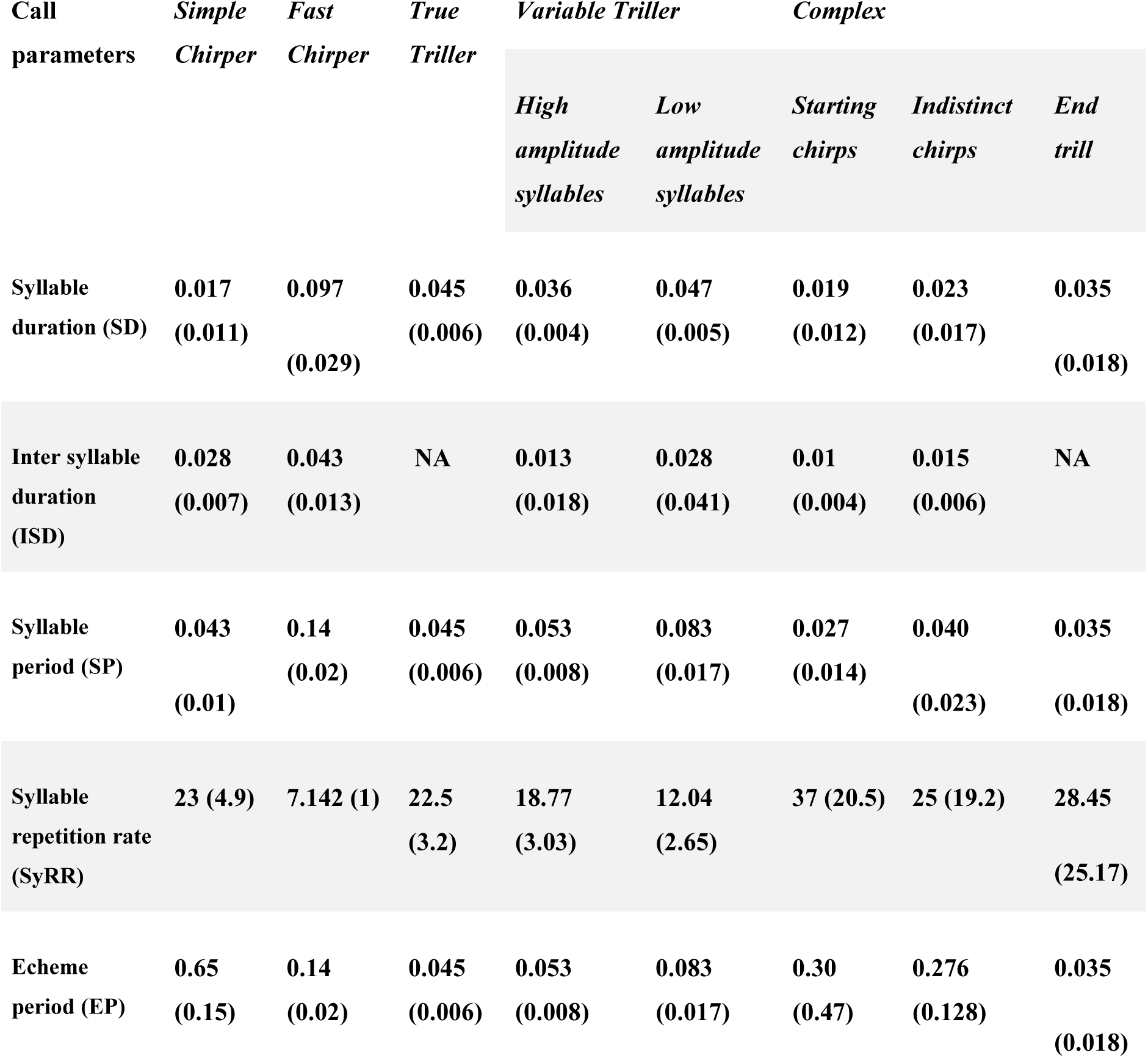
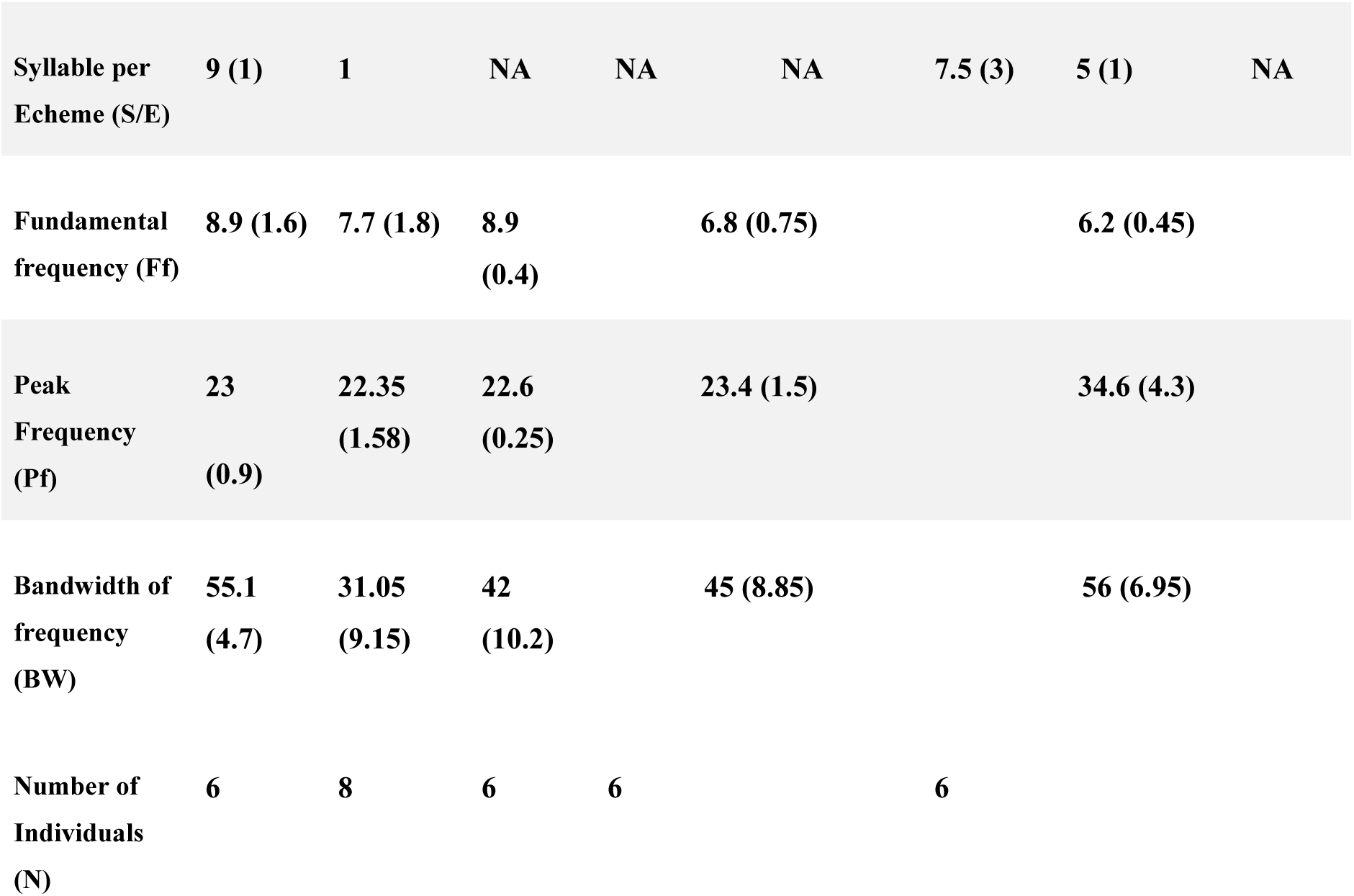
Median (+/− Inter quartile range) values of all the temporal and spectral call features quantified for the five call types of *Mecopoda* we found. Note that for all triller call types and trilling parts of calls, there is no echeme period; and the syllable duration and inter-syllable duration are not calculable for trills with indistinct syllables - in these cases syllable period is equal to syllable duration.

The first two principal components explain ∼99% of the variation between different call types (Table 2). The first principal component has contributions from moderately positive syllable duration, syllable period and fundamental frequency and moderately negative syllable reputation rate, echeme period, peak frequency, and bandwidth, while the second principal component had contributions from moderately negative syllable repetition rate and syllable duration and moderately positive contributions from the remaining parameters.

**Table 2.** Acoustic call characters and their contribution to PC1 and PC2.

| Call features | PC1 | PC2 |
| --- | --- | --- |
| Syllable duration (SD) | 0.42 | -0.08 |
| Inter syllable duration (ISD) | 0.35 | 0.2 |
| Syllable period (SP) | 0.4 | 0.09 |
| Syllable repetition rate (SyRR) | -0.3 | -0.13 |
| Fundamental frequency (Ff) | 0.2 | 0.4 |
| Peak Frequency (Pf) | -0.3 | 0.125 |
| Bandwidth of frequency (BW) | -0.4 | 0.618 |
| Echeme period (EP) | -0.156 | 061 |
| Syllable per Echeme (S/E) | -0.25 | 0.54 |
| Eigenvalue | 4.95 | 2 |
| Explained (%) | 71.24 | 28.7 |

### Fine temporal structure comparison

The syllable repetition rate emerges in the PCA as the most variable fine temporal parameter between call types. A direct comparison of syllable repetition rate amongst all the call types reveals that the SyRR of “Fast Chirper” is significantly lower than “True Triller” and “Simple Chirper” (P = 0.03, 0.02 respectively). There is no significant difference between “True Triller” and “Simple Chirper” in terms of just the syllable repetition rate, and their separation in the PCA is not related to syllable structure. “Variable Triller” shows similarities in fine temporal structure with all three call types except the “Complex” call type (p = 0.05).

### Spectral structure comparison

The scatter plot between the fundamental and peak frequencies of different call types shows that only callers of the “Complex” call type are significantly different from the other four call types. Only the “Complex” call type separates out from the other call types (Fig. 13 A - C), with a significantly higher peak frequency (p values in Table 3) and lower fundamental frequency from all the other call types (p values in Table 4). “Variable Triller” has significantly lower fundamental frequency and significantly higher peak frequency than “Simple Chirper” but overlaps with both “Fast chirper” which is variable in fundamental frequency as well as “True Triller” with no significant differences in either peak or fundamental frequency. “True Triller”, “Simple Chirper” and “Fast Chirper” have similar spectral characteristics to one another, with no mutually significant differences in fundamental or peak frequency. However, when considering the frequency bandwidth, both “Complex” and “Simple chirper” calls are different from the other three call types but similar to each other. “True Triller” and “Variable Triller” call types show no difference in bandwidth features. “Fast Chirper” shows a variable range in bandwidth (Table 5).

**Figure 13.**
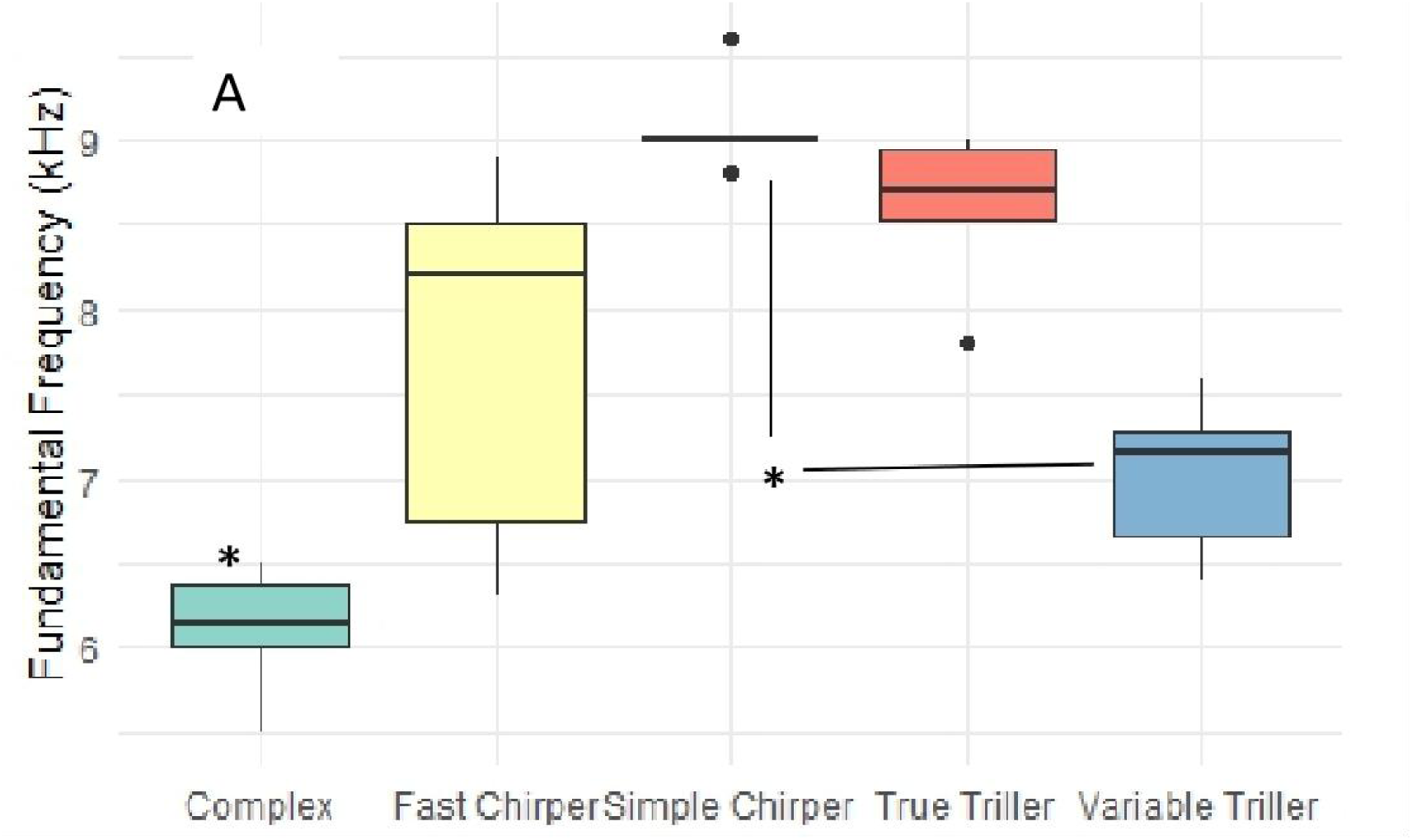
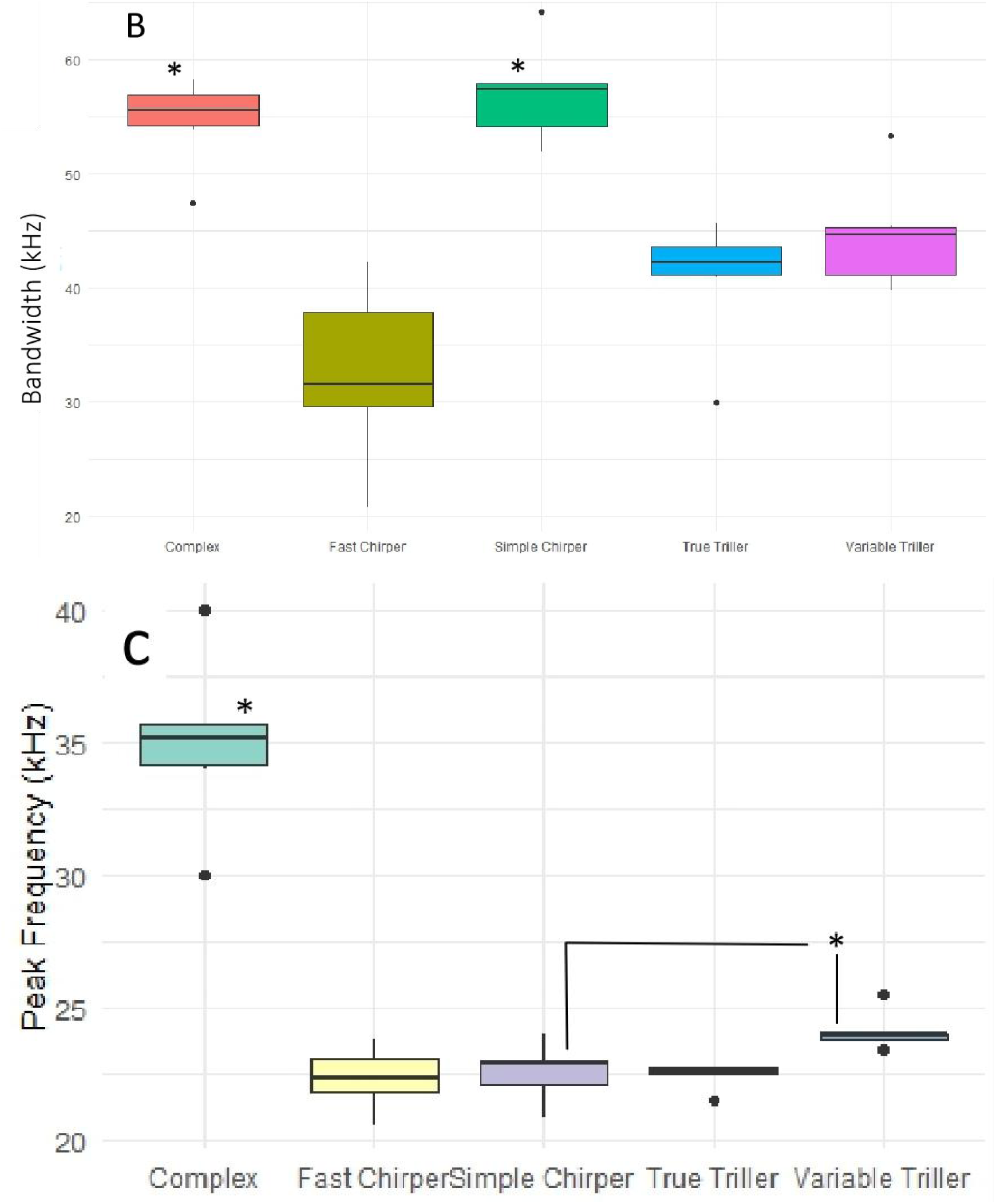
(A) Fundamental frequency; (B) frequency bandwidth; and (C) peak frequency of the five different call types of *Mecopoda*.

**Table 3.** P-values of the non-parametric paired Dunn’s test for differences in fundamental frequency between each call type.

| P-values | CO | FC | SC | TT |
| --- | --- | --- | --- | --- |
| FC | 0.01* |  |  |  |
| SC | 0* | 0.27 |  |  |
| **TT** | 0* | 1 | 1 |  |
| **VT** | 0.02* | 1 | 0.03* | 0.23 |

**Table 4.**
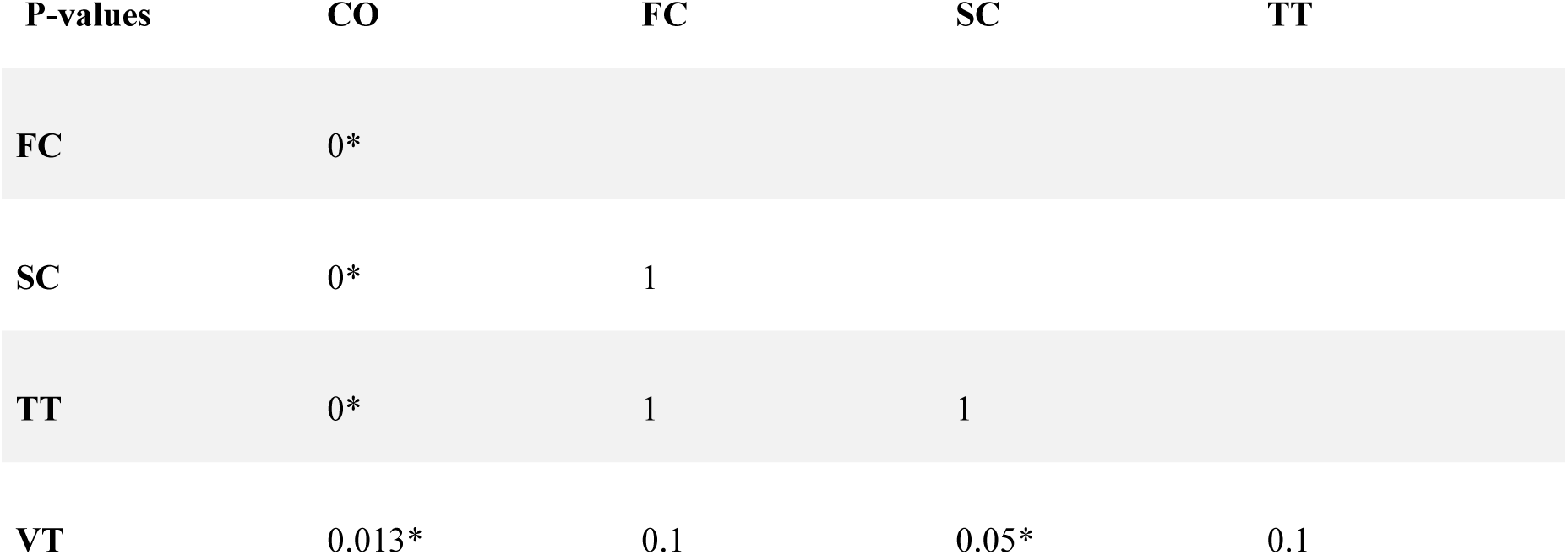
P-values of the non-parametric paired Dunn’s test for significant differences in peak frequency between each call type.

**Table 5.** P-values of non-parametric paired Dunn’s test for difference in frequency bandwidth between each call type.

| **P-Values** | **CO** | **FC** | **SC** | **TT** |
| --- | --- | --- | --- | --- |
| **FC** | 0* |  |  |  |
| **SC** | 1 | 0.002* |  |  |
| **TT** | 0.003* | 1 | 0.01* |  |
| **VT** | 0* | 0.4 | 0.01* | 1 |

### Hierarchical clustering of call types based on acoustic characters

Cluster analysis corroborates the PCA, with only the “Complex” call forming a separate cluster from the other four call types (Fig 14). Within the second cluster of four call types, “Simple Chirper” separates out as most distinct, while “Fast Chirper” and “True Triller” form separate clusters whose closest neighbors are part I and part II of the Variable triller call respectively. A couple of individual calls of “Variable Triller” are found within the “Complex” cluster, and the two parts of the “Variable Triller” call show a messy separation from each other. Likewise, different parts of the “Complex” call exhibit no specific cluster formation relative to each other.

**Figure 14.**
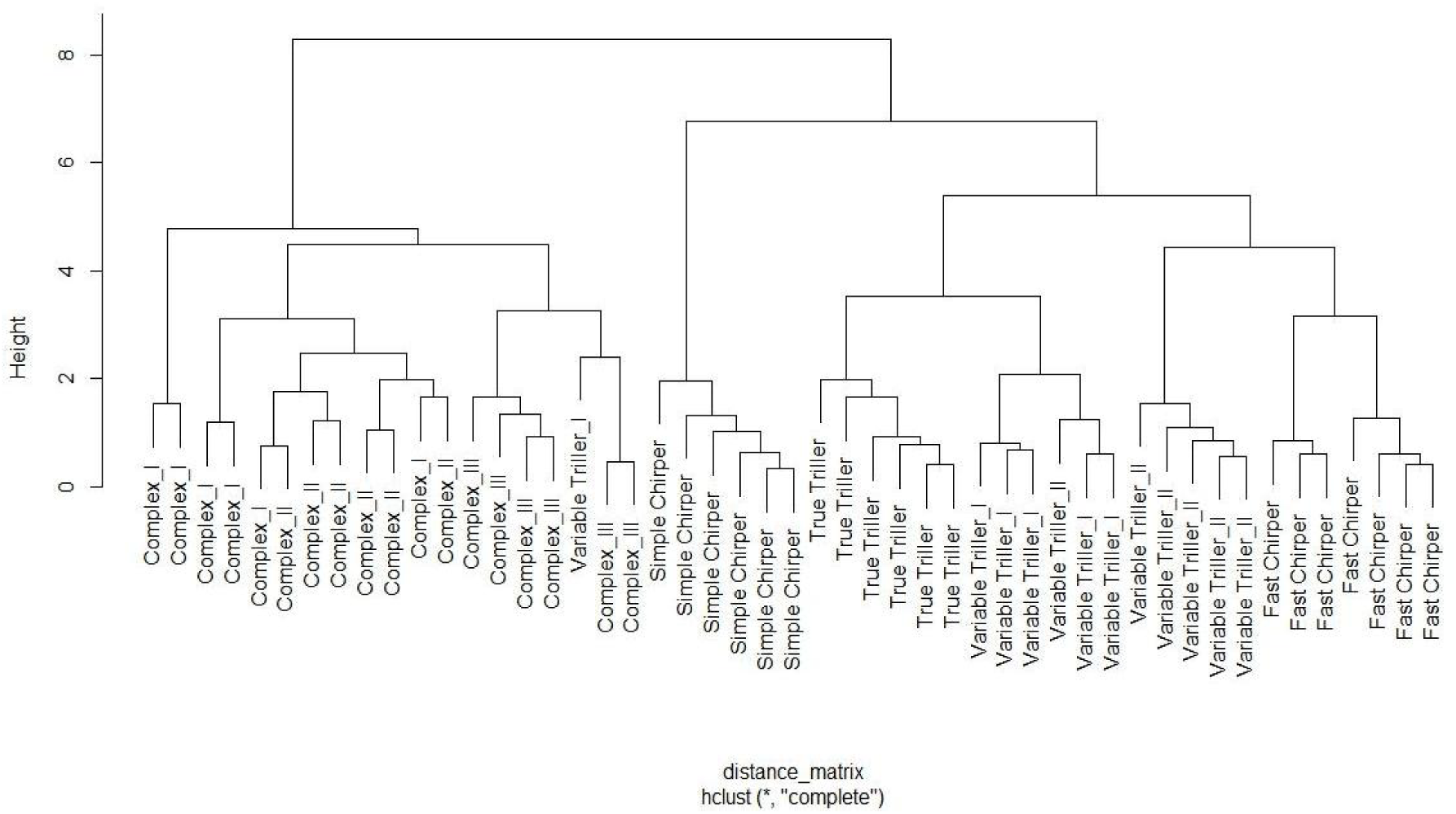
Dendrogram of different *Mecopoda* call types based on hierarchical cluster analysis of temporal and spectral characteristics of calls.

### Morphological structure analysis

All morphological characters were divided into two categories: quantifiable morphological features (i.e., body length, eye length, etc.) and qualitative morphological features (i.e., shapes of supra-anal plates).

### Quantitative feature measurement with PCA

Measurements of 21 different morphological structures were quantified and subjected to a PCA to examine any morphological feature-based separation between the different call types (Table 7). The first principal component has contributions from the forewing length of the third femur and third tibiae and the combined length of the body and wing - mostly traits relating to body sizes. The second principal component has positive contributions from the eye length and width, head length, and sternum length. Clustering across the first two principal components explained approximately 51% of the variance between different call types (Table 6), with no distinct clustering or overall quantitative morphological separation across the various individuals between the five sympatric call types of *Mecopoda* at Nokrek National Park (Fig 15).

**Figure 15.**
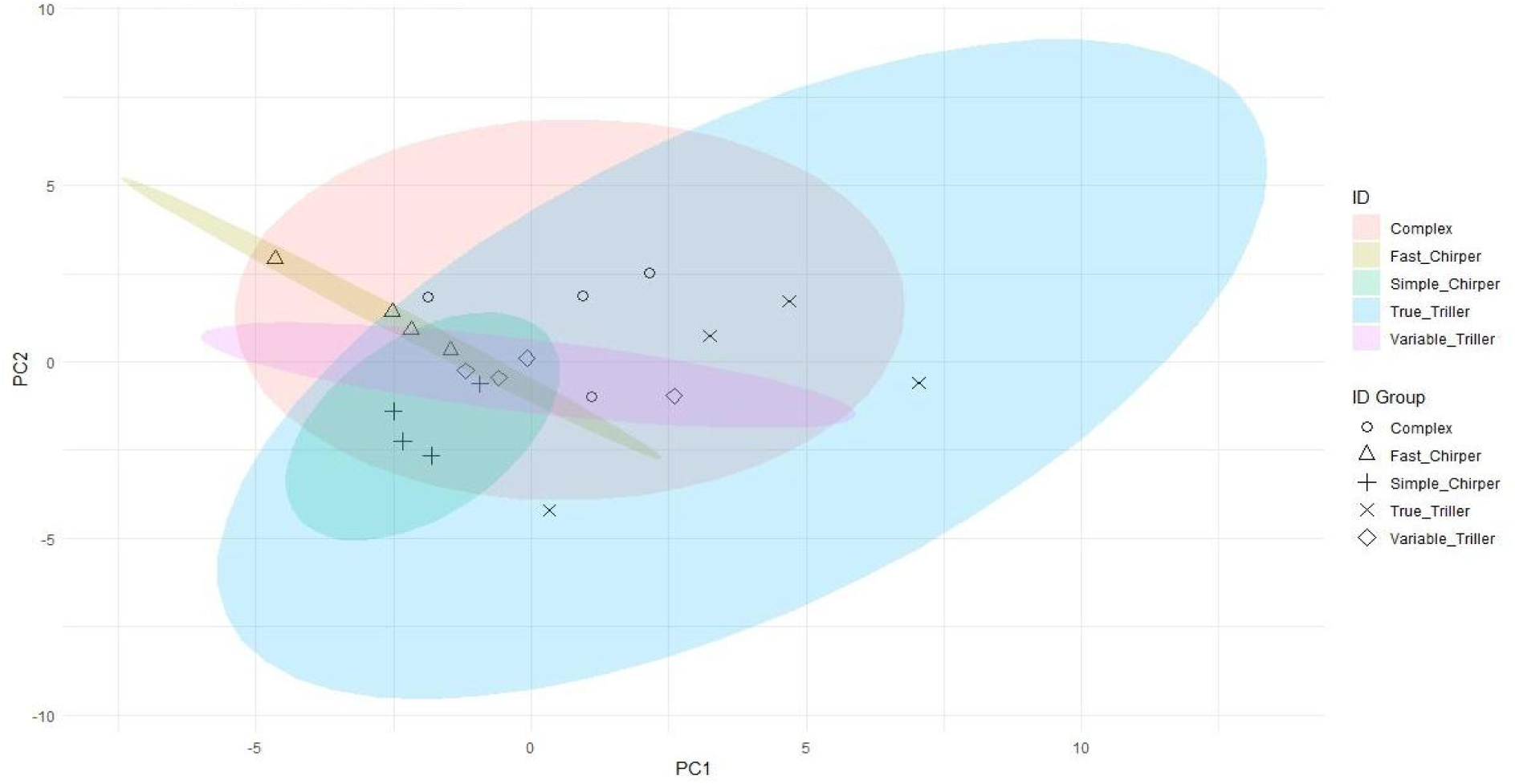
PCA visualizing the grouping of each call type of *Mecopoda* based on quantitative morphological characters. Each marker represents one individual of a particular call type, and colors denote the spread of each call type.

**Table 6.** PC1 and PC2 values of each quantitative morphological characters with eigenvalues and explained variance.

| Parameters | PC1 | PC2 |
| --- | --- | --- |
| PH | 0.107 | 0.195 |
| PL | 0.198 | 0.144 |
| HL | 0.0891 | 0.310 |
| EL | 0.033 | 0.3675 |
| EW | 0.044 | 0.213 |
| SL | 0.130 | -0.301 |
| LBA | 0.087 | 0.295 |
| CL | 0.169 | -0.467 |
| StL | 0.145 | 0.239 |
| BL | 0.170 | 0.276 |
| FWL | 0.295 | -0.051 |
| BL+WL | 0.314 | -0.0134 |
| FWW | 0.323 | -0.0359 |
| **HWL** | 0.116 | -0.433 |
| **HAW** | 0.039 | -0.157 |
| **F1** | 0.3 | 0.013 |
| **T1** | 0.313 | -0.002 |
| **F2** | 0.302 | -0.07 |
| **T2** | 0.321 | -0.106 |
| **F3** | 0.282 | -0.109 |
| **T3** | 0.237 | -0.064 |
| **Eigenvalue** | 7.98 | 3.29 |
| **Explained variance (%)** | 36.3 | 14.9 |

**Table 7.** Median values (+/− Interquartile range) of all the morphological measurements (in mm) quantified for the five call types of *Mecopoda* we found.

|  | **Morphological Parameters** | *Simple Chirper* | *Fast Chirper* | *True Triller* | *Variable Triller* | *Complex* |
| --- | --- | --- | --- | --- | --- | --- |
| 1 | **Pronotum Height (PH)** | 5.35 (0.5) | 6 (0.7) | 6.15(0.42) | 6 (0.07) | 6 (0.15) |
| 2 | **Pronotum Length (PL)** | 7.45 (0.7) | 7.8 (0.6) | 8.6 (0.62) | 8.5 (0.2) | 8.15 (0.2) |
| 3 | **Head Length (HL)** | 2.65 (0.2) | 3 (0.6) | 3 (0.35) | 2.8 (0.05) | 3.1(0.3) |
| 4 | **Eye Length (EL)** | 1.55(0.17) | 2 (0.1) | 1.75 (0.42) | 2 (0.05) | 2.1(0.05) |
| 5 | **Eye Width (EW)** | 1.3(0.23) | 1.35(0.11) | 1.35 (0.2) | 1.35 (0.29) | 1.4(0.07) |
| 6 | **Scaups Length (SL)** | 1.25(0.15) | 1 (0.22) | 1.35 (0.17) | 1.5 (0.27) | 1.1(0.07) |
| 7 | **Length between two antennae (LBA)** | 2.5(0.07) | 2.8 (0.2) | 2.7 (0.22) | 2.8 (0.1) | 2.85(0.24) |
| 8 | **Cerci Length (CL)** | 2.8(0.3) | 2.67(0.05) | 2.95 (0.3) | 3.05 (0.17) | 2.8(0.2) |
| 9 | **Sternum Length (StL)** | 7.1 (0.75) | 7(0.4) | 8.15 (1.07) | 7.6 (1.12) | 7.5 (0.6) |
| **10** | **Body Length (BL)** | 33.5 (1.5) | 35 (0.5) | 36.5 (2) | 35 (0.5) | 34 (1) |
| **11** | **Fore Wing Length (FWL)** | 53 (0.75) | 52.5 (4) | 58 (2.75) | 53.5(1.8) | 55.5 (1.75) |
| **12** | **Body with Wing length (BL+WL)** | 61.5 (1.5) | 62.5(4.25) | 68 (5.25) | 62.5 (3) | 64 (3) |
| **13** | **Fore Wing Width (FWW)** | 14 (0) | 13.5(1.25) | 17.5 (2) | 14 (1) | 15 (0.2) |
| **14** | **Harp Length (HAL)** | 11 (0.25) | 10 (0.5) | 11 (0.5) | 11 (0.25) | 10 (0.25) |
| **15** | **Harp Width (HAW)** | 5 (0.5) | 6 (1.2) | 6 (0.25) | 6 (0.25) | 5 (0.5) |
| **16** | **Fore Femur length (F1)** | 10.5 (1.25) | 10 (0.25) | 13 (0.75) | 11 (0.5) | 11.5 (1.25) |
| **17** | **Fore Tibia length (T1)** | 11 (0.25) | 10 (0) | 13 (2) | 10.5 (1.5) | 12 (0.5) |
| **18** | **Middle Femur length (F2)** | 12.5 (1.25) | 12.5 (1.5) | 15 (0.75) | 14 (0.25) | 14 (0.5) |
| **19** | **Middle Tibia length (T2)** | 13 (0.5) | 13 (0.25) | 15 (0.5) | 13.5 (1.25) | 14.5 (1.5) |
| **20** | **Hind Femur length (F3)** | 41 (4.5) | 37.5 (3.5) | 44.5 (1.7) | 41 (0.5) | 43.5 (4) |
| **21** | **Hind Tibia length (T3)** | 42.5 (7.25) | 36.5 (3.5) | 44.5 (2) | 40 (0.75) | 43 (1.75) |
|  | **Number of individuals (N)** | 4 | 4 | 4 | 4 | 4 |

### Comparison between fore wing length and whole-body length between call types

The PCA reveals that forewing and body length are the two quantitative characters that explain most of the variance between call types. A direct comparison of whole-body length including the wing reveals no significant variation between all the call types although “True triller” is non-significantly larger than the others (Fig 16A-B). When only the forewing length is considered, differences emerge. The “True triller” wing is significantly larger than that of the “Fast chirper” (P = 0.03) and “Simple chirper” call types (P = 0.04), but not significantly different from “Variable triller” and “Complex.

**Figure 16.**
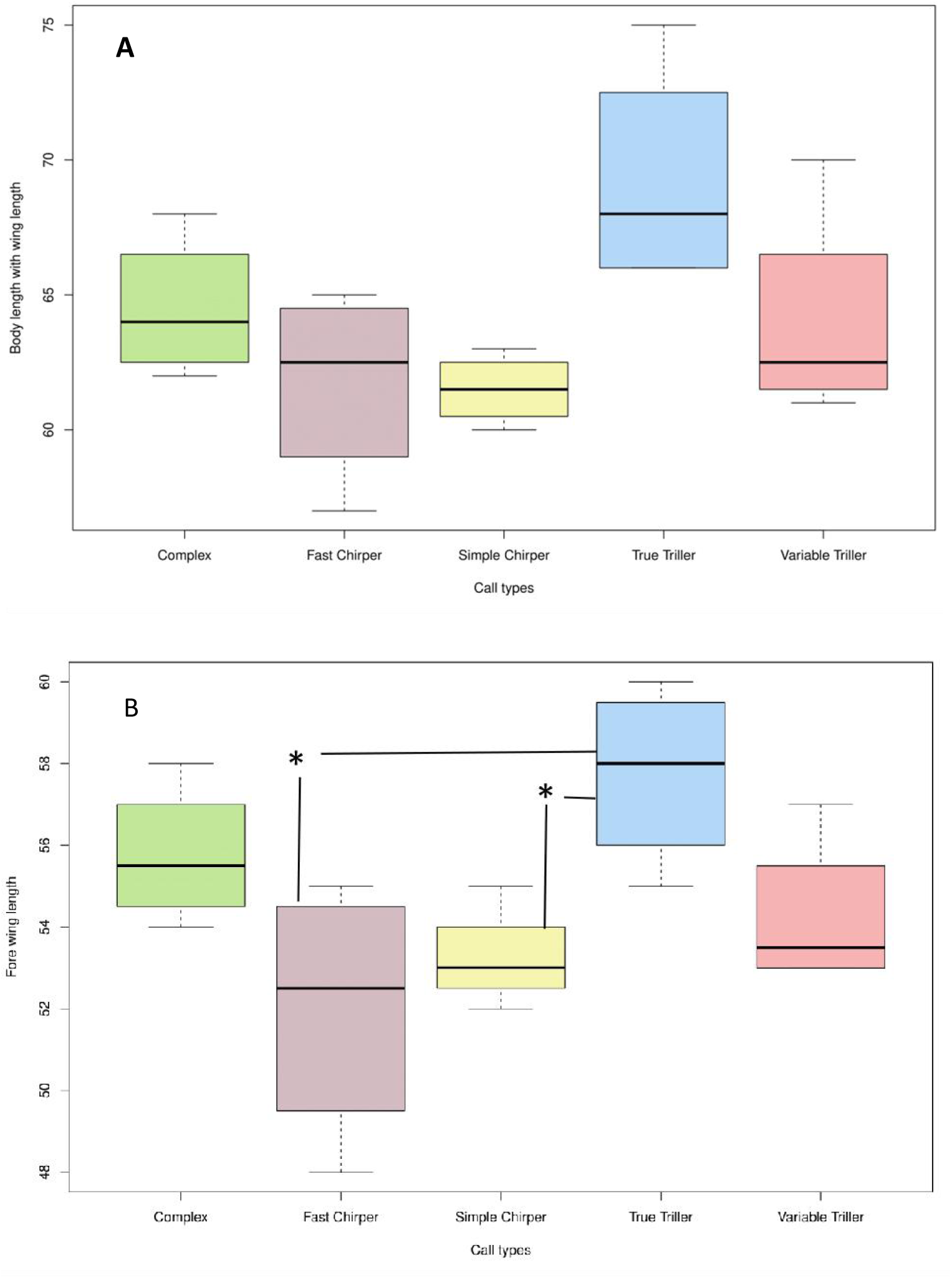
(A) Box plot showing the size of body length combining the wing length (in mm) for each call type of *Mecopoda.* (B) Box plot showing the size of fore wing (tegmina) length (in mm) for each call type of *Mecopoda*.

**Figure 17.**
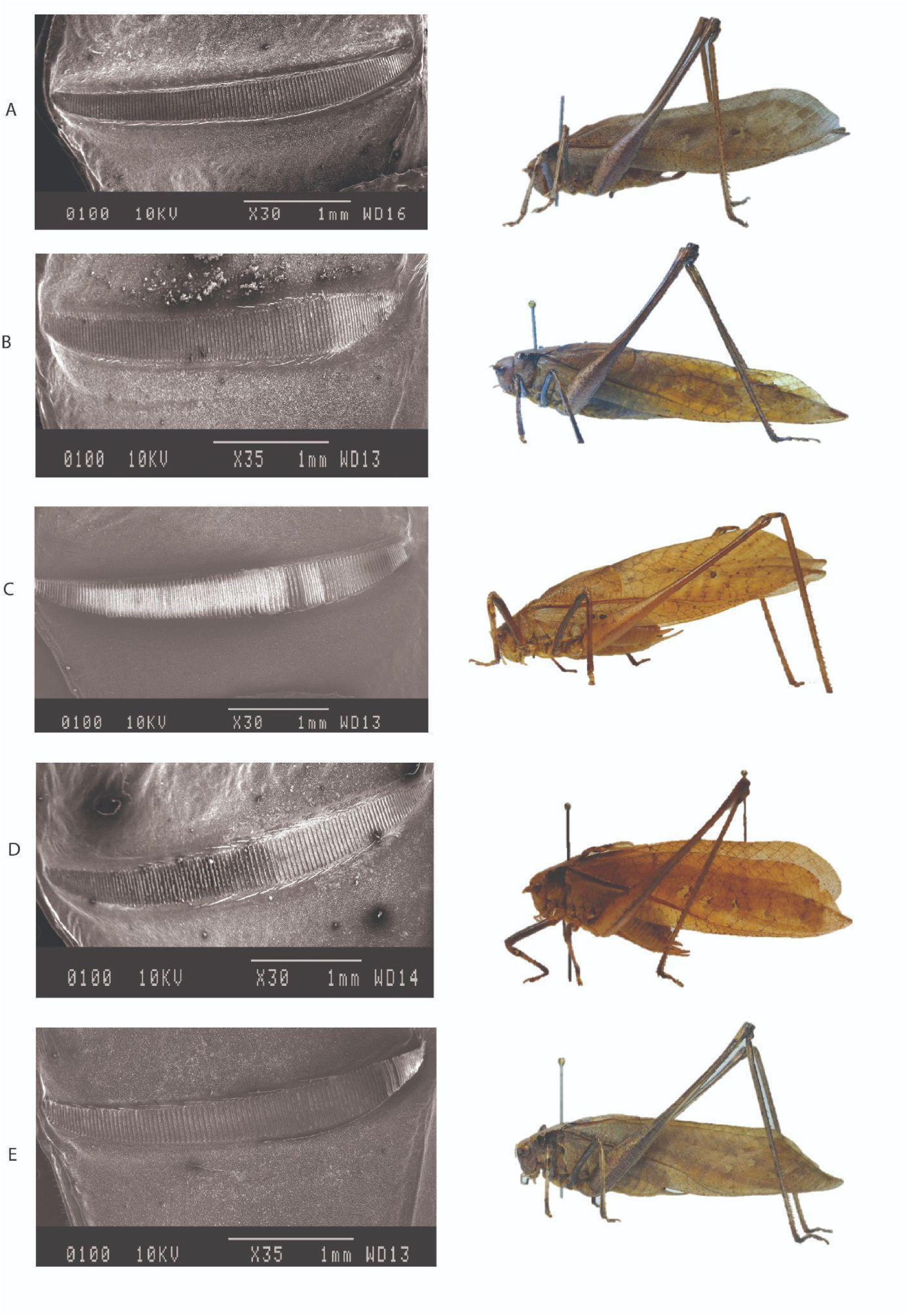
SEM images of the stridulatory file of the ventral side of left tegmen and adult males of five different call types. (A) “Simple Chirper”, (B) “Fast Chirper”, (C) “True Triller”, (D) “Variable Triller” and (E) “Complex”.

### Hierarchical clustering of all call types based on quantitative morphological characters

Hierarchical clustering based on quantitative morphological characteristics does not yield separate clusters organized by call types (Fig 18). This suggests that the call types cannot be differentiated based on quantitative morphological differences.

**Figure 18.**
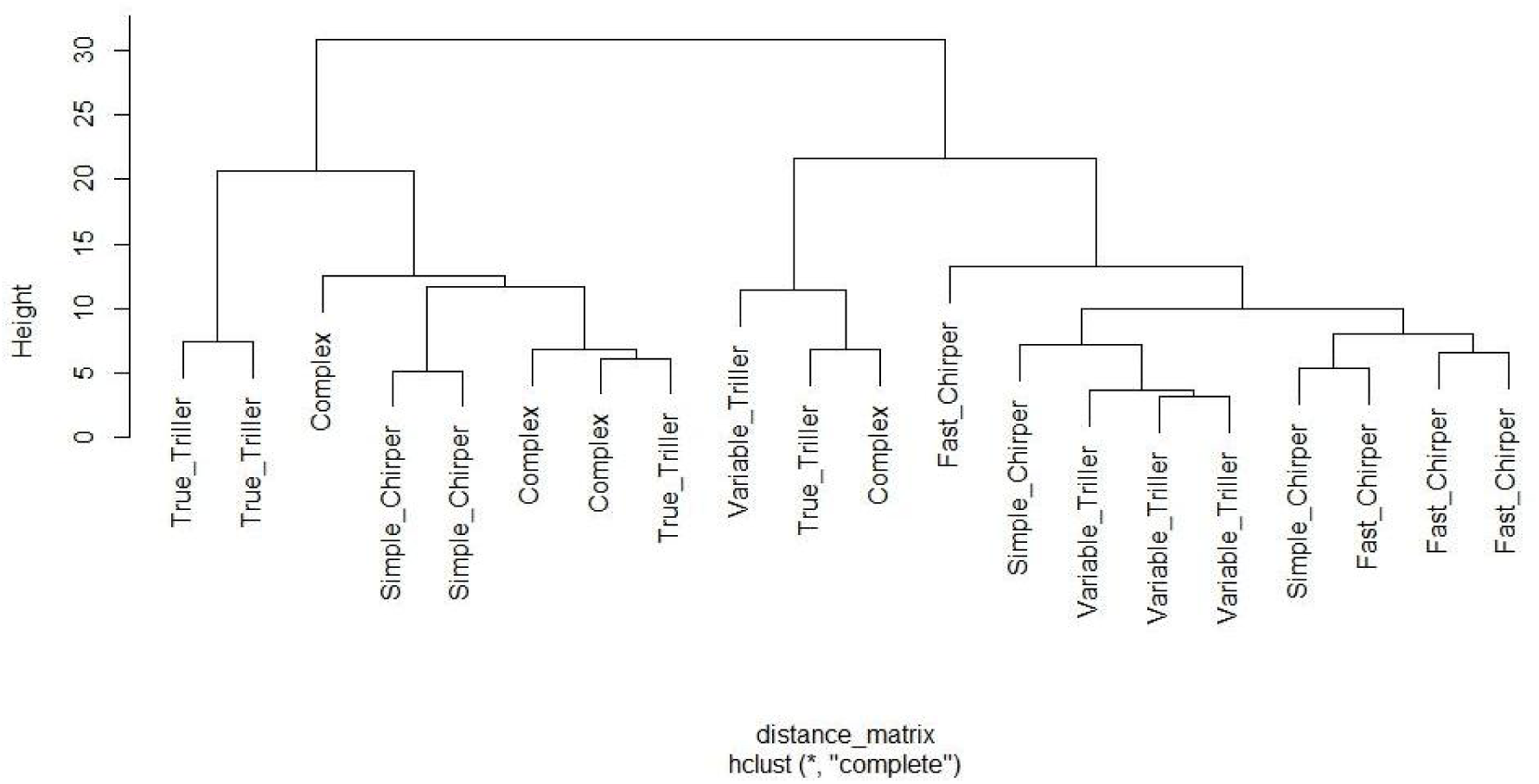
Dendrogram showing phenetic clusters of the five different *Mecopoda* call types using quantitative morphological characteristics.

### Qualitative features description

The following qualitative characters were marked across different call types: pronotum structure, stridulatory file structure, and genital morphology (Table 8). Usually, body color varies across adults from green, and grey to brown morphs, even within the same calling type (Fig 23). The Pronotum is saddle-shaped in structure, narrow in front and broad and rounded at the posterior, across individuals of different call types (Fig 19). Tegmina is elongated broad in the middle and slightly shorter than hind wings (Fig 17). Mirror cells of the right tegmen are pretzel-shaped in all *Mecopoda* call types. The file on the ventral side of the left tegmen is long and gradually narrowed in both ends in all *Mecopoda* samples (Fig 17). Sub-genital plates in all males are elongated and bifurcated into two outward anal styles. Anal cerci are stout and bent inward in the case of males of all call types (Fig 19).

**Figure 19.**
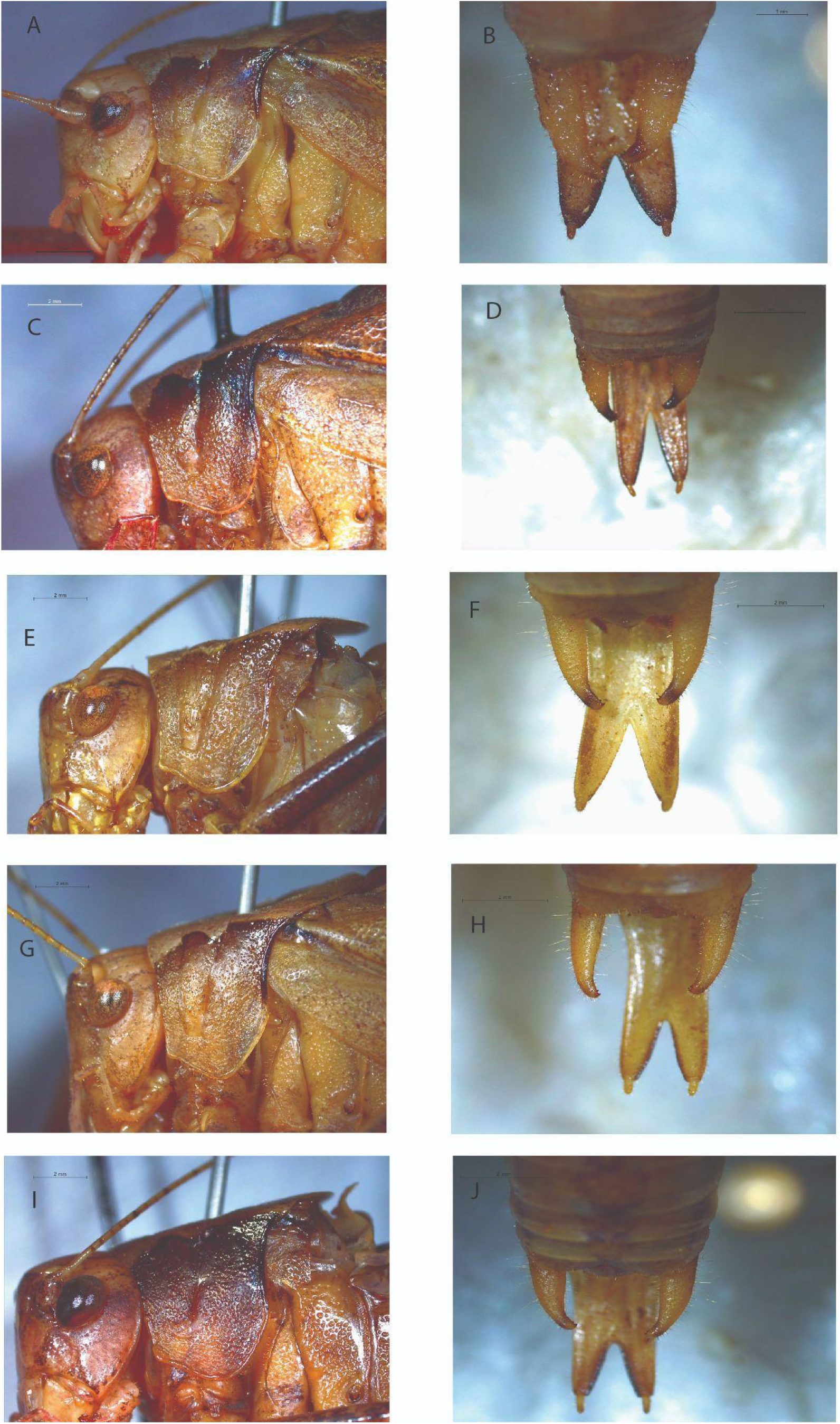
Lateral side of the pronotum and ventral side of the external male genitalia of five different call types of *Mecopoda.* (A-B) “Simple Chirper”, (C-D) “Fast Chirper”, (E-F) “True Triller”, (G-H) “Variable Triller” and (I-J) “Complex”.

**Table 8.** List of qualitative morphological characters.

| SL no | Qualitative Characters |
| --- | --- |
| 1 | Fastigium shape and structure |
| 2 | Pronotum Structure |
| 3 | Shape of Tegmina |
| 4 | Structure of file presents in ventral of left tegmen |
| 5 | Structure of mirror cells in dorsal of right tegmen |
| 6 | Structure of external genitalia |
| 7 | Structure of anal cerci |
| 8 | Structure of anal styles |
| 9 | Whole body colouration |

**Table 9.**
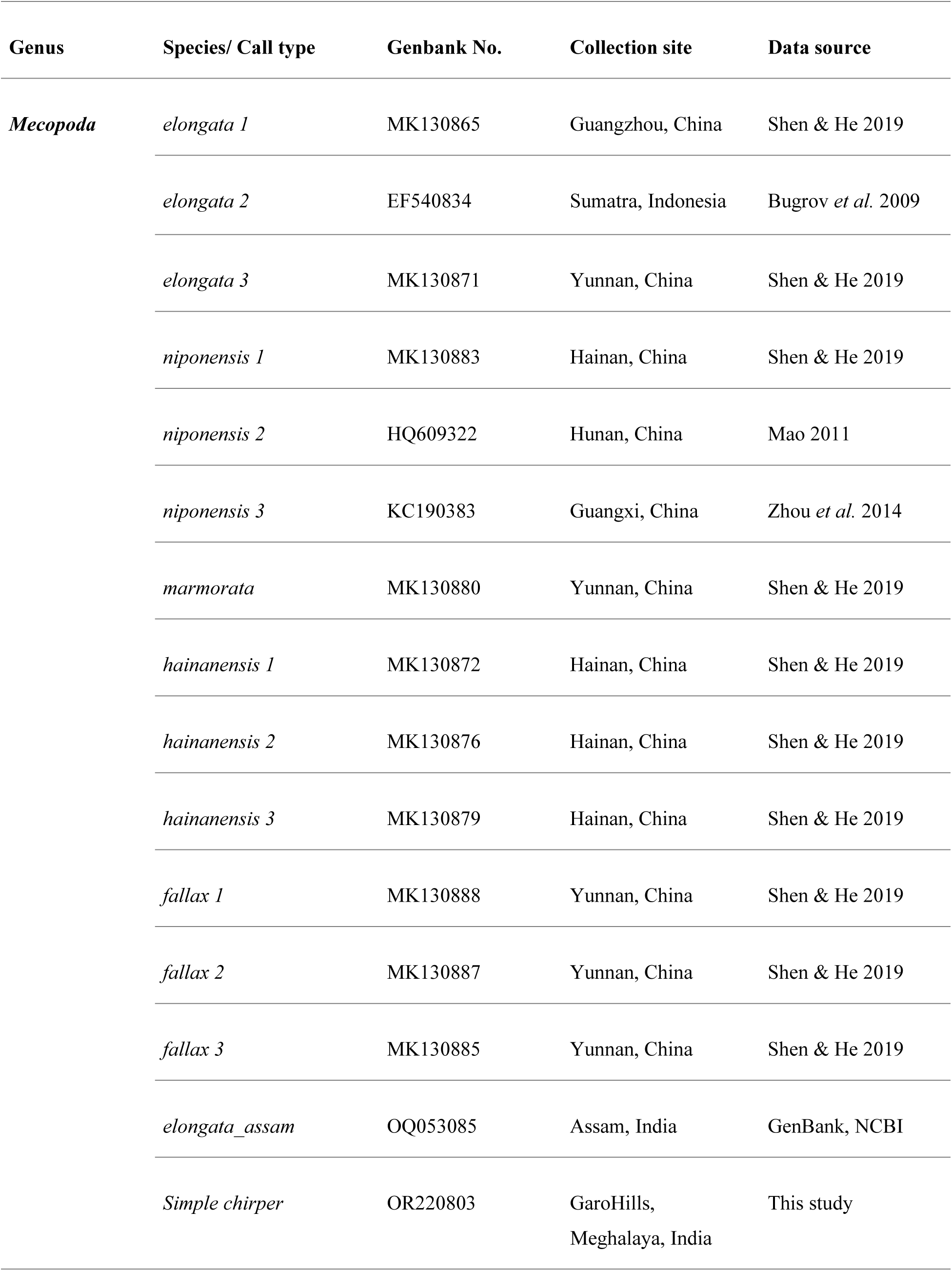
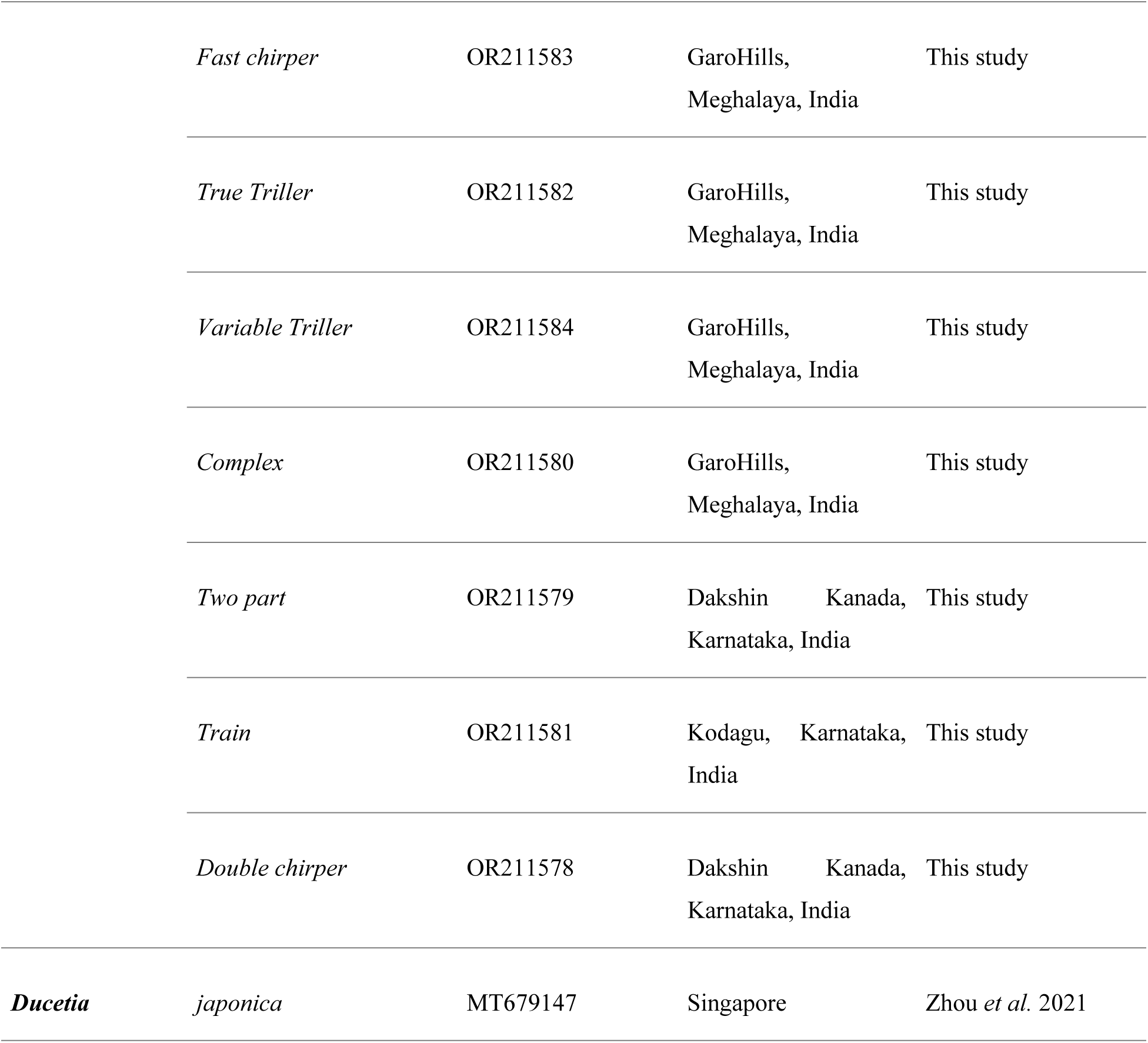
Source and collection information and GenBank accession number of all *Mecopoda* species and call types.

However, the fore wing shapes of different call types look somewhat different (Fig 21). The “Variable triller” forewing looks broader with a blunt end compared to all the chirpers call types, “Complex” and “True Triller”. Geometric morphometric analysis shows that the shapes of the forewings also show considerable inter-individual differences. Cluster analysis based on forewing shapes shows undifferentiated clusters exist across call types (Fig 20). Overall, there are no qualitative structural differences between the five call types.

**Figure 20.**
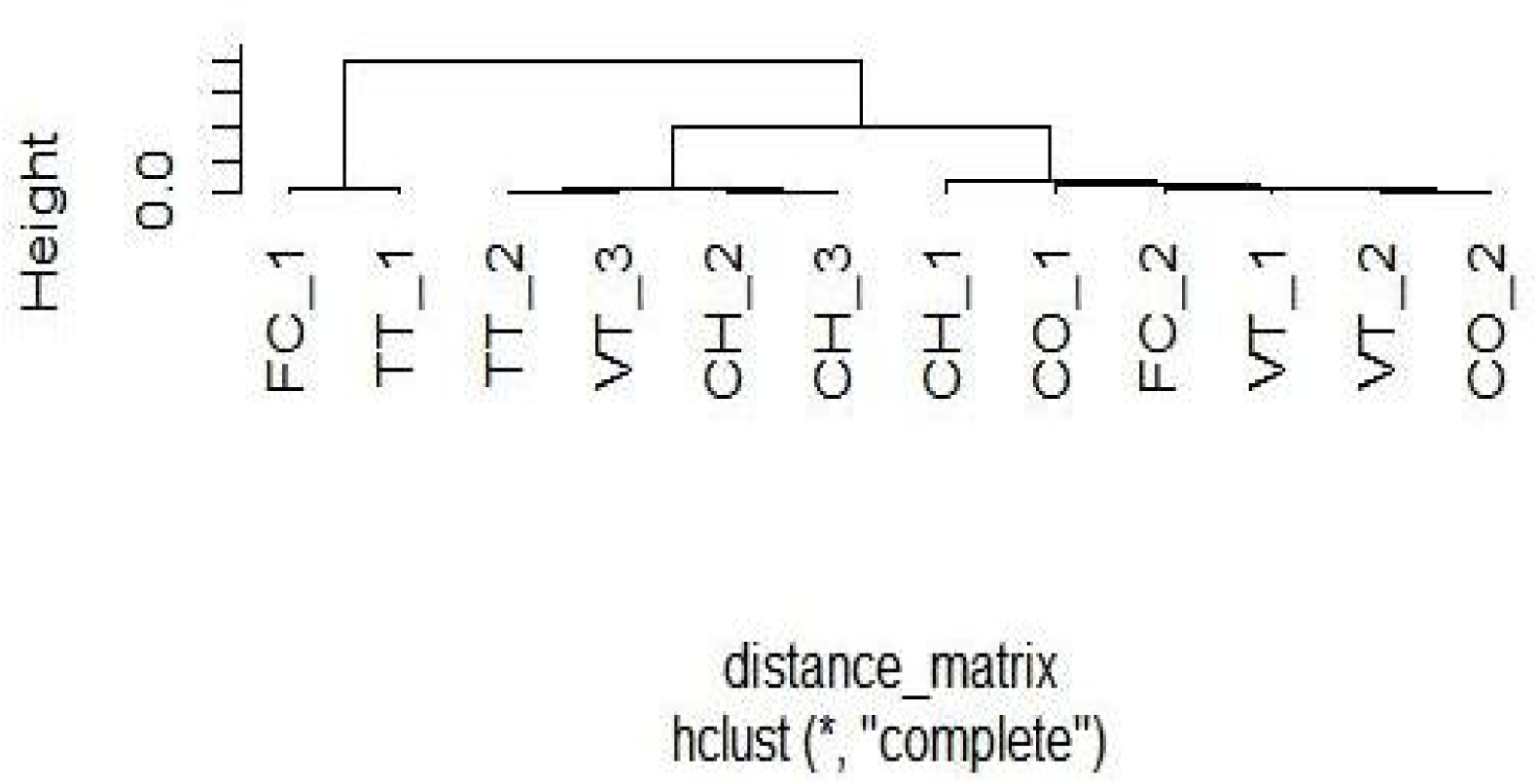
Dendrogram showing phenetic clusters of the five call types using adult tegmina shape.

**Figure 21.**
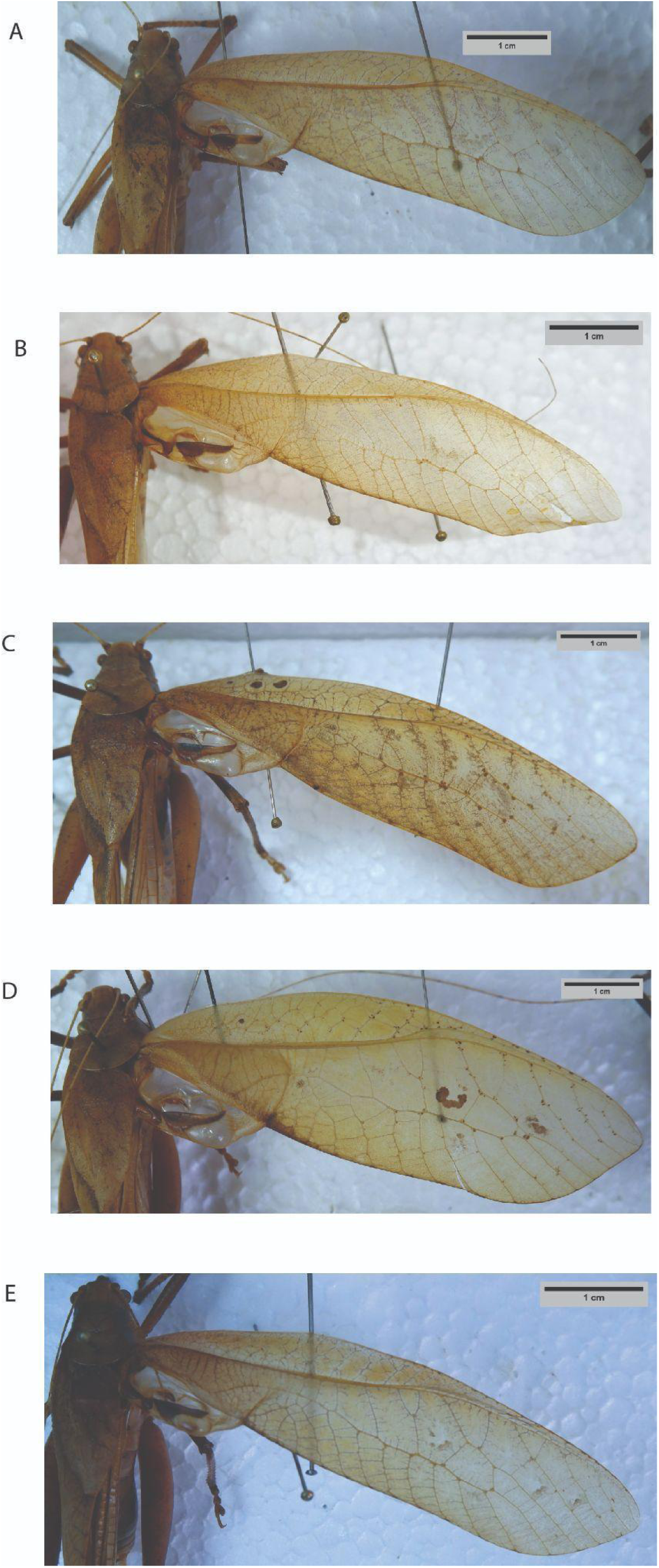
Photographic images of dorsal view of left tegmen of adult males of five different call types. (A) “Simple Chirper”, (B) “Fast Chirper”, (C) “True Triller”, (D) “Variable Triller” and (E) “Complex.”.

### Phylogenetic analysis

In this phylogenetic tree, a total of 23 COI sequences were used from various previously described *Mecopoda* species and our samples with *Ducetia Japonica* as the out-group. Accession numbers for all samples are mentioned in Table 10. The location of the collecting site of each sample is mapped out (Fig 1C). The phylogenetic tree supports the monopoly of *Mecopoda* species, within which 3 major clusters exist (Fig 22). *M. marmorata* is the only species of *Mecopoda* that is not part of any of the clusters. One cluster consists of specimens of of *M. hainanensis.* The second cluster consists of *M fallax and M niponensis,* and the third cluster consists of *M. elongata* along with various acoustic morphs of *Mecopoda* from south India. From among our 5 call types of *Mecopoda* both the triller call types, “True Triller” and “Variable Triller”, are close to the *M. fallax* cluster; while “Fast chirper”, “Simple chirper” and “Complex” call types are part of the *M. elongata* cluster along with the South Indian acoustic morphs.

**Figure 22.**
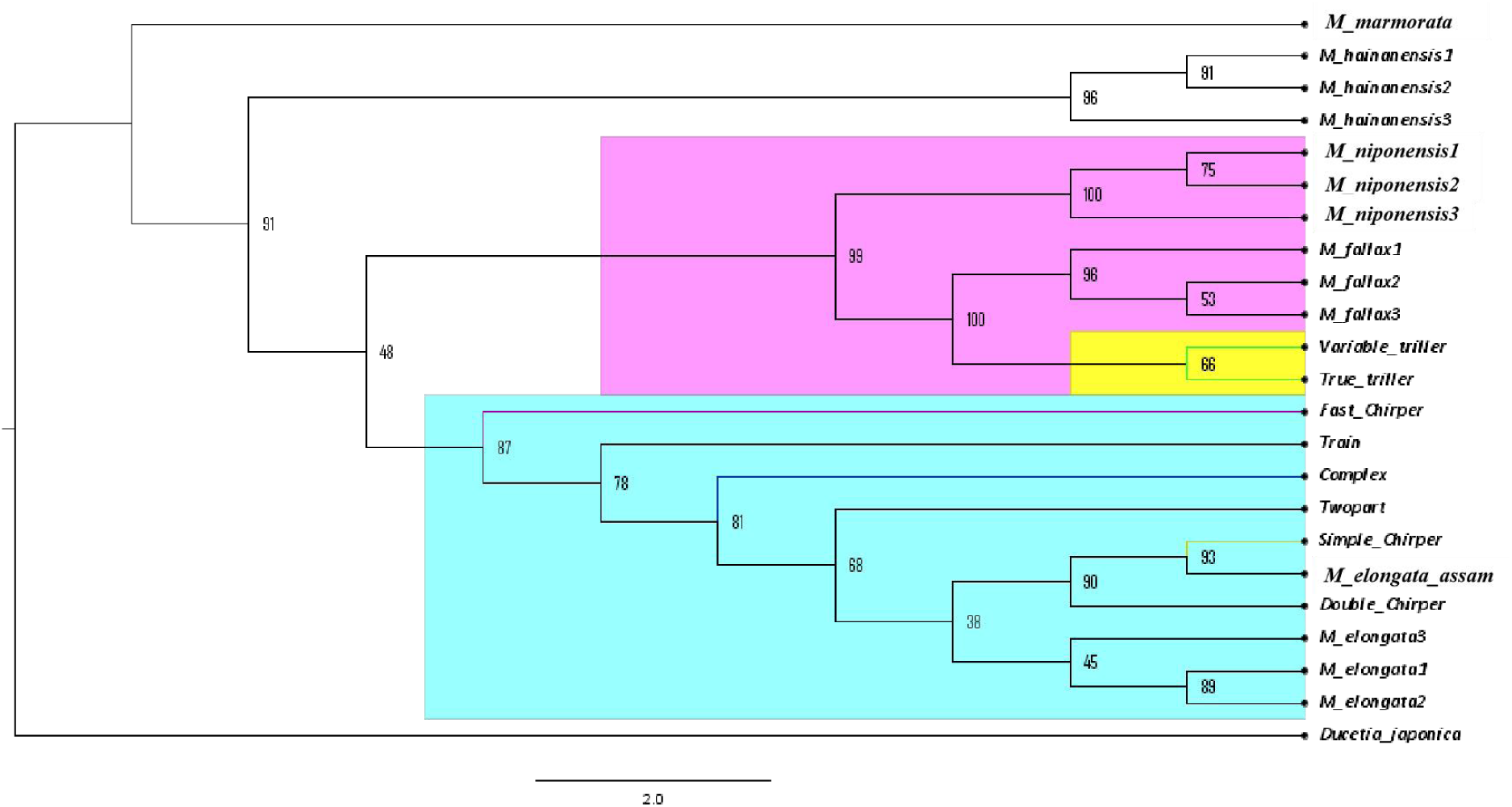
Phylogenetic tree of different *Mecopoda* species from southeast Asia and Indian call types based on COI gene. The tree was constructed via Maximum likelihood with GTR+G and was rooted with respect to *Ducetia japonica*. Bootstrap values are indicated in every node and species or call types are mentioned on the right.

**Figure 23.**
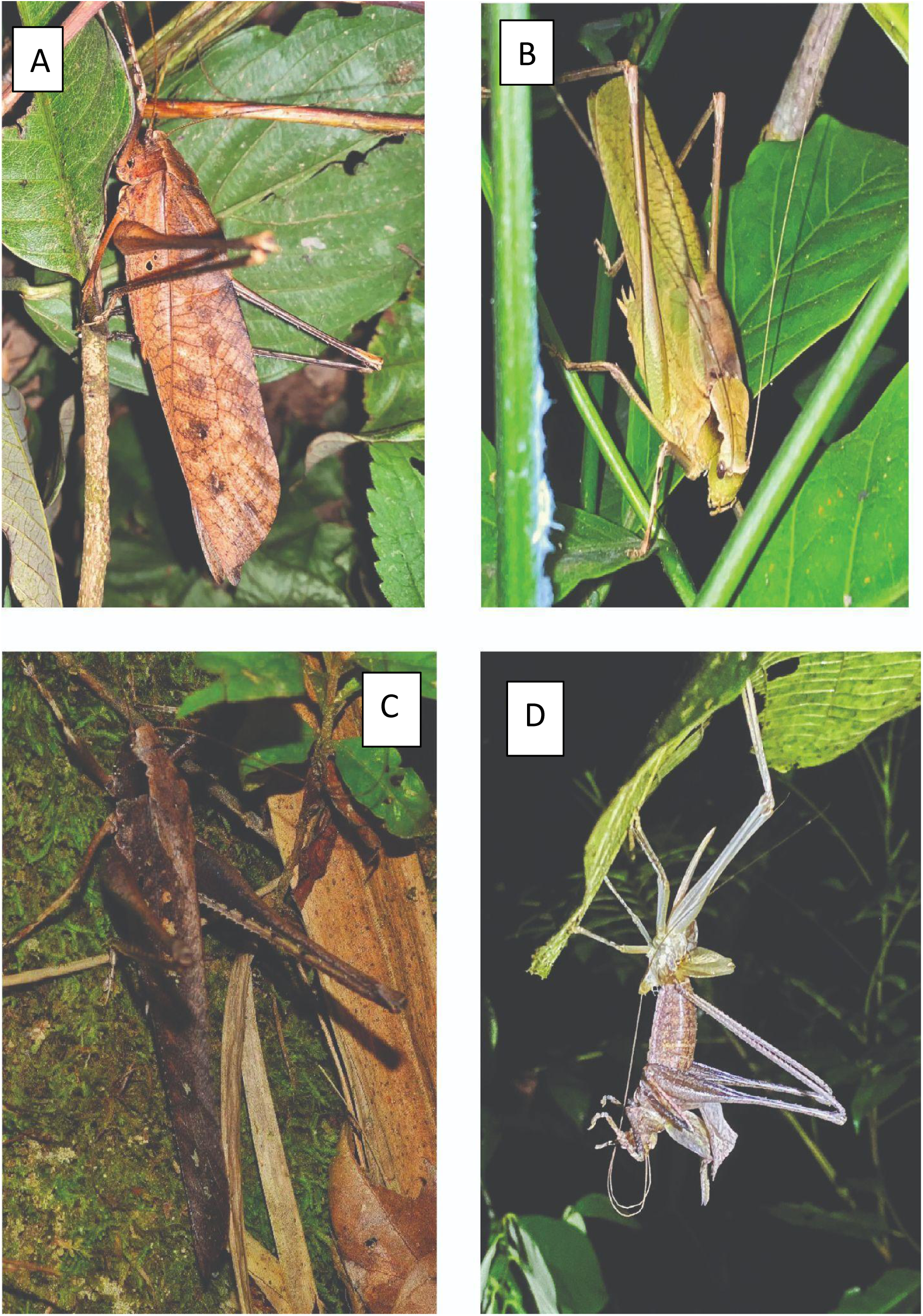
Photographs of (A, B, C) different color morphs of adult male *Mecopoda* in the wild and (D) molting of a female *Mecopoda* in the wild. All pictures were taken at Nokrek National Park, Meghalaya.

**Table 10.** Mean intra-specific distance in each species and nearest neighbor (NN) genetic distance for all call types and species.

| Species/call type | Mean intraspecific distance (%) | Nearest Neighbour (NN) | Genetic distance to NN (%) |
| --- | --- | --- | --- |
| *Mecopoda* “Simple Chirper” | N/A | *Mecopoda elongata* | 0.5 |
| *Mecopoda* “Fast Chirper” | N/A | *Mecopoda* “Train” | 4 |
| *Mecopoda* “True Triller” | N/A | *Mecopoda* “Variable triller” | 0.3 |
| *Mecopoda* “Variable triller” | N/A | *Mecopoda fallax* | 3.1 |
| *Mecopoda* “Complex” | N/A | *Mecopoda* “Two part” | 0.8 |
| *Mecopoda* “Train” | N/A | *Mecopoda* “Complex” | 1 |
| *Mecopoda* “Two part” | N/A | *Mecopoda* “Simple Chirper” | 1.8 |
| *Mecopoda* “Double Chirper” | N/A | *Mecopoda elongata_ assam* | 2 |
| *Mecopoda marmorata* | N/A | *Mecopoda hainanensis* | 6 |
| *Mecopoda elongata* | 1 | *Mecopoda* “Two part” | 2 |
| *Mecopoda niponensis* | 2 | *Mecopoda fallax* | 6 |
| *Mecopoda fallax* | 0.2 | *Mecopoda* “Fast Chirper” | 6 |
| *Mecopoda hainanensis* | 1.15 | *Mecopoda niponensis* | 7 |

The difference between our two trillers call types, “True Triller” and “Variable Triller”, is only 0.3%, but these two form a monophyletic clade that has a high genetic difference in the COI sequence from *M. fallax (*3%) and from *M. elongata* sub-group (6.5%). Our “Simple chirper” and “Complex” call types are part of a monophyletic clade along with the South Indian chirper, Two-part, Train and Double Chirper. “Simple chirper” has very low genetic differences of 1.26% from M. elongata_assam, as does Complex with a distance of 1.6% from *M. elongata*_assam. The “Fast Chirper” call type’s COI gene shows a p-distance from the neighboring “Complex” call type of 2.7%, and a distance of 3.6% with other specimens from the *M. elongata* superspecies group (Table 10), indicating comparatively higher genetic difference accumulation.

Therefore both “Variable triller” and “True triller” have considerable genetic distance in their COI gene from the rest of the *Mecopoda* species in India but have high mutual sequence similarity. “Fast chirper” also shows considerable COI gene divergence from the other call types as well as from the *Mecopoda* species from Southeast Asia for whom we have COI data. “Complex” and “Simple chirper” COI gene sequences form a monophyletic clade with the South Indian *Mecopoda* call types and *M elongata* specimens.

## Discussion

In this piece of work, we have provided a thorough acoustic and morphological characterisation of five different call types of *Mecopoda* found syntopically in the evergreen forest of Nokrek National Park in the Indo-Burma biodiversity hotspot region. We also did a phylogenetic analysis of the sequence of the single mitochondrial gene cytochrome oxidase I (COI) isolated from these call types as well as the call types previously described from south India (Nityananda & Balakrishnan, 2006).

Of these call types, our “Simple Chirper” is acoustically distinct from the “Chirper” from India described by Nityananda & Balakrishnan (2006) and other chirpers from South-East Asia described by Römer *et al*. (2002). The mean syllable period and echeme period of our “Simple Chirper” are 43 +/ 10 ms (median, +/− IQR) and 650 +/− 15 ms (median, +/− IQR) respectively, and both are higher than the Chirper syllable period of 10.4 +/− 1.1 ms (mean +/− SD) and chirp period of 483.3 +/− 43.5 ms (mean +/− SD) described previously (Nityananda & Balakrishnan., 2006).

The *Mecopoda* “triller” described by Tiwari & Diwakar (2022) from Assam sounds similar to our *Mecopoda* “Variable Triller”, except that the former has been described as having only one type of syllable with a mean syllable period and syllable duration of 19 +/− 4 ms (mean +/− SE) and 11+/−2 ms (mean +/− SE) respectively. Our “Variable Triller”, in contrast, has a varying syllable structure, based on both intensity and duration - with syllable period ranging from 53 +/−8 ms to 83 +/−17 ms (median, +/− IQR) respectively, and syllable duration from 36 +/− 4 ms to 47 +/− 5 ms (median +/− IQR). The bandwidth of “Variable triller “is 45 +/− 8.85 kHz (median +/− IQR), while the bandwidth of Mecopoda elongata “triller” is 26.83 +/− 9.57kHz (mean, +/− SE).

Our “True Triller” call type with no distinct inter-syllable interval, and a syllable period of 45 +/− 6 ms (median, +/− IQR) also differs from that *Mecopoda* “triller”. It sounds somewhat similar to *M. fallax* from Assam (Tiwari & Diwakar., 2022). However, *M. fallax* has been characterized as having a syllable duration of 15 +/− 1 ms (mean +/− SE), and an echeme duration of 30 +/− 3 ms (mean +/− SE). Since we also see two peaks within our syllable, these presumably correspond to the two syllables grouped in what they consider an echeme; however, it is unclear how the syllable period they describe of 45 +/− 64 ms (mean +/− SE) is larger and more variable than the echeme period. The repeating minimal element in the call seems to us to have a period corresponding to what we and they have both described as the syllable period. There is also a difference in bandwidth - “True triller” has a bandwidth of 42+/− 10.2 kHz (median +/− IQR) but *M. fallax* has a bandwidth of 29.1 +/− 3.9 kHz (mean +/− SE).

Our “Fast Chirper” call type sounds similar to the “Chimer” call described from Assam (Tiwari & Diwakar., 2022) but has a very distinct syllable structure. “Chimer” calls have been described as having a syllable duration of 50 +/− 1 ms (mean +/− SE) and a syllable period of 270 +/− 9 ms (mean +/− SE) while our “Fast Chirper” has a syllable duration of 97 +/− 29 ms (median +/− IQR) and syllable period of 140 +/− 20 ms (median +/− IQR). Our videos of this call type clearly show that each syllable corresponds to a single wing movement. Besides this temporal structure difference, the spectral structure varies between our “Fast Chirper” in terms of the peak frequency (median = 22.35, +/− 1.6 kHz), and bandwidth (median = 31, +/− 9.15 kHz) compared to the peak frequency (mean= 41.7, +/− 2.3 kHz) and bandwidth (median = 40.3, +/− 2.15 kHz) of *M. elongata* “Chimer”.

There is no call in the *Mecopoda* literature that seems to bear any similarity to the “Complex” call. Interestingly, “Complex” and “Simple chirper” COI gene sequences constitute a monophyletic clade along with the other South Indian *M. elongata* call types, including various other call types with verses with segments composed of chirps alternating with trills. “Fast chirper” is an outgroup to this clade, with high COI gene divergence from the other Indian call types (a p-distance from the neighboring “Complex” call type of 2.7%, and 3.6% with other specimens from the *M. elongata* superspecies group (Table 11)). “Variable triller” and “True triller” have accumulated relatively high genetic distance in their COI gene (6.5%) from the rest of the *Mecopoda* species found in India. But these two call types have high mutual sequence similarity, with a 3% genetic difference in the COI sequence from the nearest neighbor *M. fallax (*3%) and from *M. elongata* sub-group.

This can also be contextualized relative to what is known of many newly characterized cryptic species and call types of *Mecopoda* from South Asia and Southeast Asia, for whom various aspects of call acoustic structure, COI gene sequence and wing morphology have been described (Heller *et al.,* 2021; Liu *et al.,* 2019; Liu *et al.,* 2020; Gorochov *et al.,* 2020). Liu *et al* (2020) divided the *Mecopoda elongata* superspecies complex into three subgroups, or subsuperspecies (Cigliano et al., 2024), based on acoustic characteristics. *Mecopoda confracta* is one subgroup of the genus *Mecopoda* acoustically characterized by a relatively simple call structure (Heller *et al 2021*, Liu *et al 2020)*. Morphologically, species of this subgroup show the narrowest mirrors with comparatively short files and long tegmina. *M. confracta* and *M. synconfracta* have been described as distributed across China (Heller *et al*., 2021), and the Indian *Mecopoda elongata* is considered a species of this subgroup (Nityananda & Balakrishnan., 2006; Tiwari *et al.,* 2022; Ingrisch & Shishodia., 2000; Bailey., 1990; Sismondo., 1990; Rentz *et al.,* 2006). Another subgroup of the *Mecopoda elongata* superspecies is *Mecopoda elongata minor,* found in China, Korea, and Japan, defined by mixed acoustic components - discontinuous chirps at the beginning followed by a trill component (Liu *et al* 2019, Heller *et al,* 2021). Despite our “Complex” call types and the “Train” call type from the Western Ghats described by Nityananda & Balakrishnan (2006) having these complicated call characteristics, our COI gene data provide a tentative basis for considering our “Complex”, “Simple Chirper” and “Fast chirper” call types from Meghalaya, as well as the other call types from South India first described by Nityananda & Balakrishnan (2006) that we sequenced, viz. “Train”, “Double Chirper” and “Two Part”, as part of the *Mecopoda confracta* subgroup of *Mecopoda elongata*.

The other major subgroup, *Mecopoda niponenesis* includes subspecies characterized by their wider tegmina and mirror and trilling call types (Liu *et al.,* 2020) as compared to the other species of this subgroup, viz., *M fallax,* which possess a comparatively longer tegmen (Liu *et al.,* 2019). Our two triller call types “Variable Triller” and “True Triller” have *M. fallax* of China as their closest neighbor when looking just at the gene COI sequence. The wing size and shape of “Variable Triller” is also similar to *M. fallax* of China, and longer than the other three *Mecopoda* call types found in Meghalaya, although this difference is not statistically significant.

While whole genome data would be required to be certain about this grouping, with single mitochondrial COI gene data being the only genetic information available across *Mecopoda* lineages in South and Southeast Asia in order to construct a phylogenetic tree (Liu *et al*., 2019; Heller *et al*., 2021; Liu *et al*.,2020). Since COI is a mitochondrial gene with no introns and a haploid inheritance mode, *COI* is subject to far less recombination compared to the nuclear genome and is considered a reliable and valid tool for species identification (Hebert *et al.,* 2009; Saccone *et al.,* 1999). Morphological and acoustic analysis alone cannot be definitive indicators of speciation for morphologically cryptic species of the kind described in *Mecopoda* (Nityananda & Balakrishnan., 2006; Liu *et al*.,2020), especially when there are variations in colouration often found within each call type (Heller *et al*., 2021). Phenotypic plasticity subject to the organism’s life stage (Wilson-Wilde *et al.,* 2010) or as a response to environmental cues (Fusco & Minelli., 2010), may also affect morphological or acoustic traits. Only genomic data, however, will prove conclusive in terms of examining the relationship between call types and species identity in this genus that shows high call divergence and cryptic speciation.

Apart from Nityananda & Balakrishnan (2006) and Tiwari & Diwakar (2022), there are a few other descriptions of *Mecopoda* species from the Indian subcontinent. Walker (1869) described *Mecopoda pallida* from North India (Liu et al., 2020), and Aswathanaryana & Aswatha (1994) described *Mecopoda sp.* based on karyotype data, but they provide no acoustic information to compare with our findings. So far, all known acoustic call types have been taxonomically described as *Mecopoda elongata* (Linnaeus., 1758; Karny., 1924), and Heller et al. (2021) has described the Indian call types as endemic on the basis of their acoustic uniqueness, but as a taxonomic “black spot” in the absence of genetic data. This is the first study from India with any genetic sequencing information on the various *Mecopoda* call types from two rainforest systems in the Western Ghats of South India and Meghalaya in the North-East, in addition to acoustic and morphological data.

All five of our specimens are sympatric and syntopic, co-occuring in the same geographic location and habitat, coexisting on the same temporal and spatial scales. No geographical isolation is present between these call types, but they are geographically well separated from the call types described from the Western Ghats in South India (Nityananda & Balakrishnan, 2006) and separated by much smaller distances from the call types described by Tiwari & Diwakar (2022). It is worth noting that we did not find most of these call types, other than the “Simple Chirper” type in other parts of Meghalaya, from our field station in Umling to Shillong, the capital of Meghalaya, to the Byrdaw falls at the southernmost edge of Meghalaya. There is therefore no continuous distribution of these call types across Meghalaya, let alone contiguous with the gibbon sanctuary in Assam where Tiwari & Diwakar (2022) did their fieldwork. Likewise, apart from “Chirper”, the call types from the Western Ghats have not been described from locations outside the forests contiguous with the Ghats (Nityananda & Balakrishnan, 2006). However, it is interesting that this pattern of call type divergence in *Mecopoda* is found within two major rainforests of India, the Western Ghats montane rainforests and the subtropical rainforests of Meghalaya in the Indo-Burma hotspot.

It is common for populations from different geographic locations to accumulate genetic divergence, and divergence in terms of acoustic and morphological characters (Jang & Gerhardt., 2005, Ghosh et al., 2023). However, in the case of sympatric populations, especially when synoptic, genetic and morphological divergence is expected to accumulate between subgroups only when some form of reproductive isolation emerges. Divergence in calling songs can start to bring about assortative mating, especially when divergent call choice preferences emerge among mating partners (Ritchie., 1996, Wagner & Reiser., 2000). This can eventually have a feedback effect of furthering call divergence (Higashi *et al.,* 1999), leading to increasing behavioral reproductive isolation (Hill *et al.,* 1972; Bhattacharya *et al*., 2017), and even speciation (Higashi *et al.,* 1999). If genetic divergence accumulates in the absence of morphological divergence, this can give rise to cryptic species (Jones., 1997), where genetic divergence accumulates to the point of making the process of crossing over during meiosis highly error prone, rendering offspring of mating across genetically diverged types infertile (Orr., 1995), in alignment with the traditional biological species concept (Myar., 1963). Walker *et al.,* (2003) proposed that every call type in sympatry may in fact represent a different species (Shaw., 1990). Since call structure in crickets appears to be a genetically encoded, unclear behavior (Hoy., 1974), independent mutations leading to variations in call structure and call preference will only lead to speciation if they initially lead to assortative mating. Assortative mutation would result in reproductive isolation and speciation only without substantial crossbreeding between different variants before they accumulate large genomic differences over the generations. Mendelson & Shaw (2002) observed that the species boundary is very porous in the case of sympatric species such as *Laupala sp*., and in fact, interbreeding between two major call types may also increase the variety of sympatric call types in the population, as previously reported among katydids (Ritchie., 1996).

Out of our five five call types, except for the “Complex” call type, the other four call types show some level of overlap in terms of spectral features. In this, “Complex” is relatively unique since the Western Ghats call types were largely found to be spectrally statistically indistinguishable. The “Complex” call type shows low genetic distance from the other chirper call types, but not the trillers despite having trilling components in its song. “Variable Triller” shows some overlap in temporal acoustic features with “True Triller”; otherwise, all our call types form separate clusters based on their acoustic characteristics. We see no cluster separation based on qualitative or quantitative morphological characteristics. Wing morphology in particular is often correlated with calling songs of closely related acoustically diverging subspecies due to the role of wings in call production (Pitchers *et al*, 2014), and genital morphology may also mediate reproductive isolation. However, in our case, neither genital nor wing morphological data show significant differences across call types. However, there are other traits whose divergence might contribute to reproductive isolation. Cuticular hydrocarbons have also been characterized for some of the call types of sympatrically diverging *M. elogata* from the Western Ghats (Dutta *et al.,* 2018). The key question whose answer is only partially answered is the question of the specificity of female mate choice - at least one population of one call type of Mecopoda *sp* from South India was found to show no preference between different call types (Dutta *et al.,* 2017). The difficulty with ascertaining preferences across call types is that females have been reluctant to show phonotactic behaviour within the lab, and with their cryptic morphology, silent females are impossible to distinguish based on call type. Future work on whole genome analysis of *Mecopoda* call types is needed to fully understand the biogeography and evolution of this species. *Mecopoda* also constitutes an excellent model system to look at acoustic divergence, sexual selection, speciation and sensory evolution.

## Acknowledgements

We thank the State Forest Departments of Meghalaya and Karnataka for permission to conduct fieldwork in protected areas. We are grateful to Dr. Sudipta Tung’s lab and Dr. Imroze Khan’s lab for sharing lab space and microscopy equipment. We thank Dr. Shivani Krishna, Dr. Manjari Jain, and Dr. Ranjana Jaiswara for academic guidance. We are thankful to Dr. Meghna Agarwala for helping us with the shape files for QGIS, and Dr. Kritika Garg and Dr. Balaji Chattopadhyay for guiding us in phylogenetic analysis. We also extend gratitude to our field guide Mr. Salim Sangma for their contribution and help in field collection. We acknowledge the immense support from the A’chik community people from Garo in Meghalaya during our field stay. We also extend our gratitude to our colleagues at the Biology, Psychology, and Environmental Studies departments at Ashoka University.

